# Ecological and Genetic Determinants of Essential Oil Diversity in Mediterranean *Thymus*

**DOI:** 10.64898/2025.12.09.693220

**Authors:** José C. del Valle, Esther María Martín-Carretié, Francisco José García-Cárdenas, David Doblas, Mª Ángeles Ortiz, John Thompson, Perrine Gauthier, Bodil K. Ehlers, Diego Nieto-Lugilde, Regina Berjano

**Author notes:** Correspondence: Regina Berjano.

## Abstract

The Mediterranean Basin is a hotspot of plant diversity, with many species producing aromatic essential oils (EOs) that mediate ecological interactions and stress responses. Within Lamiaceae, the genus *Thymus* shows remarkable chemical variability, yet high intraspecific EO variation often limits its taxonomic resolution. We investigated EO composition, genetic structure, and environmental influences across 39 populations of *Thymus* sect. *Mastichina* in the Iberian Peninsula, using GC-MS alongside soil and climatic data to assess drivers of chemical variation and refine taxonomic characterization. We identified 14 major EO compounds, dominated by oxygenated monoterpenes. Most populations exhibited 1,8-cineole-rich chemotypes, yet seven populations showed linalool dominance, and multiple chemotypes often co-occurred within the same populations, revealing high intrapopulation chemical diversity. Minor compounds, including camphor, borneol, and camphene, varied among genetic clusters and were significantly correlated with temperature and precipitation gradients. Differences in EO composition were also detected between ploidy levels and genetic groups, although the major compounds (1,8-cineole and linalool) remained relatively consistent, indicating both conserved and locally adaptive chemical traits. These findings suggest that EO diversity in *Thymus* sect. *Mastichina* arises from a complex interplay of environmental conditions, genetic background, and ploidy. Integrating chemical, genetic, and ecological data provides a robust framework for understanding the evolutionary and ecological drivers of EO variation in Mediterranean aromatic plants, with implications for taxonomy, conservation, and the study of adaptive chemical traits.

## Introduction

The Mediterranean Basin harbors an exceptional wealth of plant diversity, representing roughly 10% of the world’s vascular flora (Greuter 1991; Cowling *et al*. 1996; Myers *et al*. 2000; Thompson 2005; Blondel *et al*. 2010; Nieto Feliner *et al*. 2023). This region is also remarkable for its abundance of aromatic taxa, with nearly half of the recorded genera including species that produce fragrant essential oils (EOs) (Ross & Sombrero 1991; Thompson *et al*. 2003). These aromatic compounds may play multiple beneficial roles, such as attracting pollinators (Knudsen & Tollsten 1993; Zhang *et al*. 2023), defending plants against herbivory and pathogens (Valladares *et al*. 2002; Solórzano-Santos & Miranda-Novales 2012), mitigating abiotic stress (Mansinhos *et al*. 2024), or signaling neighboring plants to activate defenses (Heil & Karban 2010), among others. Such ecological versatility, along with other morphological adaptations for stress tolerance, has likely contributed to the diversification and success of aromatic lineages in the Mediterranean, a region characterized by a stressful climate with hot, dry summers and cool, wet winters (Mooney *et al*. 1974; Cowling *et al*. 1996; Thompson 2005). In addition to these ecological functions, EO composition is often explored for its potential taxonomic value. Yet, despite reflecting ecological differentiation, their frequent high intraspecific variability limits their utility as diagnostic characters (Thompson *et al*. 2003; Keefover-Ring *et al*. 2009). Nevertheless, when combined with genetic analyses, EO profiles can enhance resolution and support more accurate identification of closely related species (Salgueiro *et al*. 2000; Tzakou & Constantinidis 2005; Da Silva *et al*. 2017; György *et al*. 2020; Etri & Pluhár 2024).

The family Lamiaceae is characterized by its chemical richness, showing a remarkable diversity of EOs across different species (Thompson *et al*. 2007; Matesanz & Valladares 2014; Aedo *et al*. 2017). Within this family, the genus *Thymus* stands out as one of the most diverse and chemically variable groups, being an important component of the Mediterranean shrubland flora (Morales 1986, 1997; Melendo *et al*. 2003). Studies of this genus have revealed substantial variability in the chemical composition of their EOs, even at the infraspecific level. For instance, the wild thyme (*T. vulgaris*) displays seven distinct EO chemotypes, each defined by its predominant monoterpene (Keefover-Ring *et al*. 2009).

More recently, Kryvtsova *et al*. (2022) documented that EO composition can vary with microhabitat conditions even within the same species. Similarly, Wester *et al*. (2020) identified up to seven distinct chemotypes of *T. pulegioides* within a single limestone grassland, further highlighting the chemical complexity of this genus.

The Iberian Peninsula, a major center of diversification for *Thymus*, provides an ideal framework to further explore chemical diversity within this genus and to assess the potential taxonomic relevance of EO composition. The genus *Thymus* L. is represented by approximately 250 species classified into eight sections mainly distributed throughout the Mediterranean and Eurasia regions (Jalas 1971; Morales 1986, 2002). The section *Mastichina* Mill. is endemic to the Iberian Peninsula and traditionally includes two gynodioecious species: the endangered *T. albicans* Hoffmanns. & Link, mainly confined to the southwestern region, and *T. mastichina* L., which is widely distributed across most of the Iberian Peninsula (Moreno 2008; Morales 2010; Buira *et al*. 2017). The latter species is further divided into two subspecies that differ in ploidy and distribution: the tetraploid *T. mastichina* subsp. *mastichina* L., broadly distributed across the region, and the diploid *T. mastichina* subsp. *donyanae* R. Morales, restricted to the sandy soils of Doñana National Park (Huelva, Spain) and the Algarve (Portugal) (Girón *et al*. 2012). Recent phylogenomic analyses, however, have revealed the existence of two main evolutionary lineages that only partially align with the currently accepted taxonomic framework. One linage largely corresponds to the tetraploid *T. mastichina* subsp. *mastichina*, and the other encompasses diploid populations traditionally assigned to *T. mastichina* subsp. *donyanae* and *T. albicans*, along with some populations identified as *T. mastichina* subsp. *mastichina* (García-Cárdenas et al., in rev.). A closer examination of the diploid lineage reveals the existence of multiple geographically differentiated genetic clusters, with most populations showing evidence of genetic introgression both among diploid groups and between diploids and tetraploids (García-Cárdenas et al., in rev.).

In *Thymus*, chemotypes frequently fail to align with taxonomic or genetic boundaries, as EO profiles are highly variable at local and infraspecific scales and often shaped by environmental factors. For instance, the EO composition in *T. vulgaris* from southern France exhibit well-defined spatial patterns that are largely shaped by climatic and edaphic gradients (Gouyon *et al*. 1986; Salgueiro *et al*. 1997; Thompson *et al*. 1998, 2003, 2007; Delgado *et al*. 2014), and Tunisian populations of *T. capitatus* also exhibit marked differences in EO composition across contrasting bioclimatic zones (Jaouadi *et al*. 2018).

Within section *Mastichina*, studies of Portuguese populations revealed chemical differences among species, primarily in the contents of 1,8-cineole and linalool (Salgueiro *et al*. 1997; Miguel *et al*. 2004; Figueiredo *et al*. 2008). However, most of the chemical variability seems to occur at the infraspecific level, with populations near the Atlantic coast showing notably higher linalool content than inland populations, a pattern similarly reported in other related Iberian taxa (e.g., *T. carnosus* and *T. villosus*) (Salgueiro *et al*. 1995, 1997, 2000). Such spatially structured variability highlights the potential influence of local environmental factors in shaping the biosynthesis and relative proportions of terpenoids.

This spatial and infraspecific variability in EO composition in the section *Mastichina* suggests that local environmental factors might play an important role in shaping chemical profiles. However, such patterns also raise the question of whether some of the observed differences may reflect underlying genetic differentiation among populations (Bataillon *et al*. 2022). In this study, we aim to integrate genetic, chemical, and ecological data to refine the characterization of *Thymus* sect. *Mastichina*. Since polyploids often diverge from diploids in their morphological, chemical, and physiological characteristics, having higher rates of multivariate niche differentiation (Otto & Whitton 2000; Hull-Sanders *et al*. 2009; Baniaga *et al*. 2020), we expect that tetraploid *T. mastichina* displays a broader ecological niche breadth and greater chemical variability in EO composition than diploid lineages. In addition, we hypothesize that different genetic clusters might occupy distinct ecological niches and exhibit divergent chemical compositions. Finally, we assess whether the observed chemical profiles and ecological niches are primarily shaped by underlying genetic differences or by environmental conditions.

## Materials and Methods

### Plant material

Samples were collected from 39 populations across the Iberian Peninsula, covering the distribution range of *Thymus* sect. *Mastichina*, with special emphasis on the southwestern Iberian Peninsula, which harbors the highest concentration of diploid genetic groups (**Figure 1**; **Table S1**). For the biochemical analysis, we collected 4–5 leaves per plant from the middle section of a non-flowering stem showing no signs of herbivory. We sampled six hermaphroditic and six female individuals per population for chemical analyses. Sampling was conducted between May and June 2023, coinciding with the flowering period of the species. Individuals were randomly selected in the field, ensuring a similar phenological stage to minimize seasonal variability. To reduce the likelihood of sampling close relatives or resampling the same genet, individuals were spaced at least 5 meters apart. Leaves were preserved in 1 mL of pure ethanol to isolate their EOs. Samples were stored at 5°C until chromatography-based analysis.

**Figure 1.**
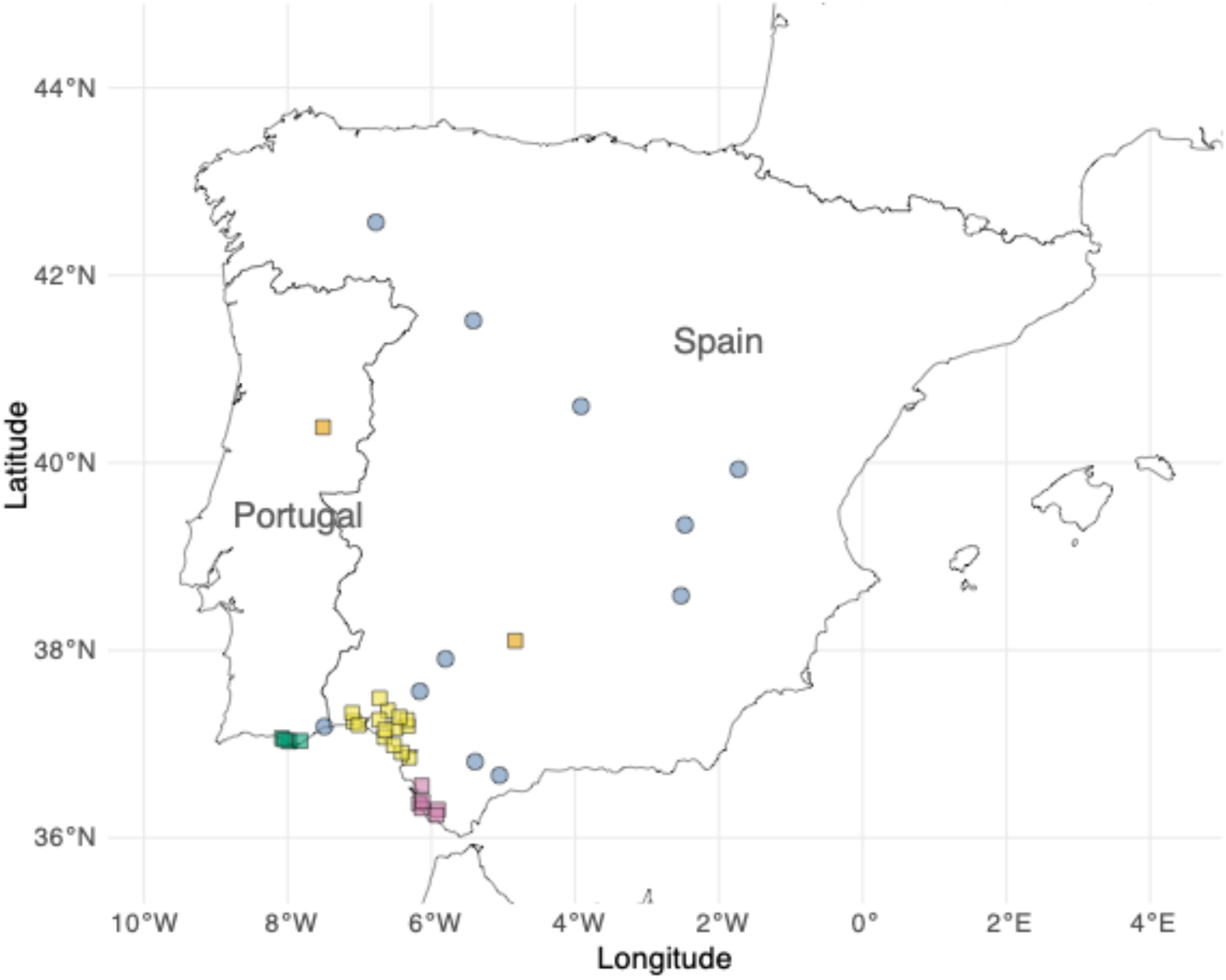
Geographic distribution of the studied populations from *Thymus* sect. *Mastichina*. The populations are color-coded according to the five genetic groups identified by García-Cárdenas et al. (in rev.): ‘*Tetraploid’* (blue), ‘*Hercynian’* (orange), ‘*Algarve’* (green), ‘*Doñana’* (yellow), and “*Cadiz*” (pink). Circles and squares represent diploid and tetraploid locations, respectively.

### Chemical analyses

Essential oils from *Thymus* sect. *Mastichina* were isolated from leaves preserved in pure ethanol. For each population, two separate pools of alcoholic leaf extracts were prepared: one from five hermaphroditic individuals and another from five female individuals, and these pools were used to generate their respective chromatographic profiles. Specifically, 4 μL of extract from each individual was combined, totaling 20 μL per pool, and then mixed with 980 μL of pure methanol to obtain a final volume of 1 mL used for chromatographic analysis. In addition, EOs from one hermaphroditic and one female individual per population were analyzed separately. Therefore, for each population and sexual morph, we obtained one individual profile and one pooled profile based on five individuals.

For chromatographic determination, a qualitative analysis of the EOs was performed using gas chromatography coupled with mass spectrometry (GC-MS). The analysis was conducted with a TSQ8000 Evo Triple Quadrupole GC-MS/MS chromatograph equipped with a ZB-5ms capillary column (30 m × 0.25 mm × 0.25 µm, Phenomenex). The injection volume was 1 μL. A general acquisition method was employed, with an MS transfer line temperature of 280.0 °C and an ion source temperature of 250.0 °C. Ionization was carried out in electron impact (EI) mode with positive polarity. The temperature program consisted of three ramps: the initial temperature was set to 50.0 °C and increased at a rate of 5.0 °C min⁻¹ until reaching 110.0 °C (first ramp). The second ramp increased at 10.0 °C min^-1^ up to 180.0 °C, followed by a third ramp at 20.0 °C min^-1^ up to 280.0 °C, which was held for 6 minutes. The injector temperature was 250.0 °C. Chromatographic data were analyzed using Thermo Scientific™ Xcalibur™ v.4.3 software (Thermo Fisher Scientific, 2016).

Identification of the components was made based on their retention times, m/z ratios of molecular ions, and ion fragmentation patterns. Additionally, the identified components were confirmed by comparison with the National Institute of Standards and Technology (NIST) mass spectrometry library integrated into Thermo Scientific™ Xcalibur™, and by matching their mass spectra with literature data on EOs previously identified in individuals of *Thymus* sect. *Mastichina* (Salgueiro *et al*. 1997; Figueiredo *et al*. 2008). The quantification of identified components was made based on GC peak areas, which were normalized relative to the total peak areas to obtain the relative proportions of each compound, enabling comparisons across pools, populations, and taxa. Some peaks were excluded as they were identified as hydrocarbon chains considered degradation products of other compounds.

Aromatic compounds were grouped into two functional categories based on their chemical nature, monoterpene hydrocarbons and oxygenated monoterpenes, and were also analyzed individually. Chemical profiling of the pooled samples revealed the presence of distinct chemotypes: while most populations exhibited a dominant 1,8-cineole-rich profile, 13 hermaphroditic and/or female pools showed higher levels of linalool than typically observed in that chemotype (i.e., linalool concentrations > 10%; see **Results**). To qualitatively determine whether the linalool peak observed at the pool level was driven by a single individual with exceptionally high accumulation or by moderate accumulation across multiple individuals within the pool, we performed additional GC–MS runs in which the five individuals making up each pool were analyzed separately, when possible.

### Environmental data

From each population, we recorded the latitude and longitude coordinates, which were then used to extract the extended bioclimatic variables (Bioclim+) from the Chelsa Climate Database v.2.1 (Brun *et al*. 2022). Specifically, we obtained values for 61 bioclimatic variables. Additionally, we collected three soil samples from each population by extracting ∼1kg of soil at 10-15 cm depth. For each soil sample, we measured standard soil-related variables following Papuga *et al*. (2018). First, soils were dried at 40°C for 48 h and stored in a cool room prior to analysis. After mixing 10 g of dry soil with 20 ml of distilled water, we separated phases using a centrifuge (10 min) and measured values in the supernatant at room temperature (c. 20°C). The pH and the electrical conductivity (c), expressed in milli-siemens per centimeter (mS/cm), were then recorded. Water retention potential (WRP) was calculated, as the percentage of water lost after drying a moist soil for 48 h at 40°C. Water retention capacity (WRC) was then calculated as the percentage of water remaining in this soil previously dried at 40°C, after drying the sample at 110°C for 5 h. Finally, organic matter (OM) was estimated as the percentage of matter lost after burning a dried sample at 500°C for 5 h.

### Statistical analyses

For each of the EOs identified, we used MANOVAs, followed by Tukey’s post hoc tests where appropriate, to assess differences between ploidy levels (diploids vs. tetraploids) and among the five genetic groups identified within *Thymus* sect. *Mastichina*: the “*tetraploid*” group and the “*Algarve*”, “*Cadiz*”, “*Doñana*” and “*Hercinian*” diploid groups (García-Cárdenas et al., in prep). The potential ecological drivers of compound-specific variation were assessed using Spearman correlations (soil and bioclimatic variables did not meet normality assumptions) with Bonferroni correction (Rice 1989) to evaluate associations between aromatic compound profiles and key environmental variables. Specifically, we included all soil-related variables considered in this study (i.e., pH, conductivity, WRP, WRC, and OM) along with eight climatic variables: bio1 (annual mean temperature), bio5 (maximum temperature of warmest month), bio10 (mean temperature of warmest quarter), bio12 (annual precipitation), bio14 (precipitation of driest month), bio17 (precipitation of driest quarter), mean relative humidity (MRH), and the aridity index (AI). These variables were chosen based on prior evidence of their influence on chemical composition in the components of *Thymus* sect. *Mastichina* (Salgueiro *et al*. 1995, 1997) and other *Thymus* species (Thompson *et al*. 2013; Llorens-Molina & Vacas 2017). Correlations were assessed at both population and individual levels. Population-level analyses used pooled data, whereas individual-level analyses included both the separately analyzed individuals and those composing the pooled samples. When no within-pool variation was detected, biochemical composition was assumed to be homogeneous in all five individuals from the pool; otherwise, the relative abundance of each compound per individual was considered.

## Results

### Essential oil composition and intrapopulation variability

When analyzing all pools from each population, we confidently identified 14 chemical compounds in the leaves of taxa from *Thymus* sect. *Mastichina*, with most compounds detected at concentrations comparable to those reported in previous studies (e.g., (Salgueiro *et al*. 1997; Miguel *et al*. 2004; Figueiredo *et al*. 2008; Delgado *et al*. 2014; Filipe *et al*. 2017; Rodrigues *et al*. 2020) (**Figure 2A**; **Table 1**). Based on their chemical nature, the compounds were classified into two functional groups: monoterpene hydrocarbons (compounds 1–6), and oxygenated monoterpenes (compounds 7–14). Monoterpene hydrocarbons were consistently present at much lower amounts than oxygenated monoterpenes, representing only 10.60% ± 3.46 (mean ± SD) of the total composition. Conversely, oxygenated monoterpenes represented the dominant class across all individuals, accounting for an average of 89.40% ± 3.46 of the total essential oil content (**Table 1**).

**Figure 2.**
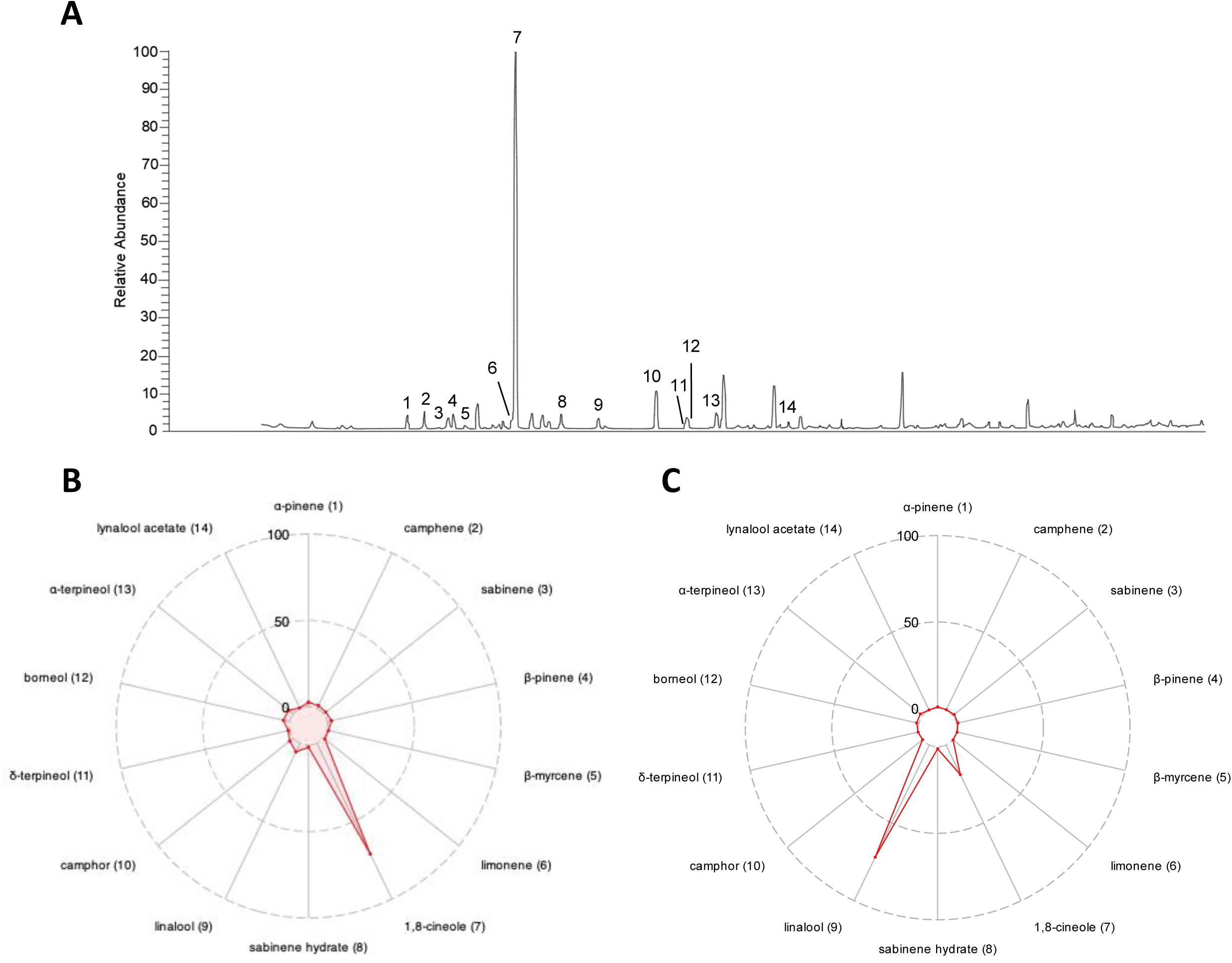
**(A)** Example of a chromatogram of leaf extract obtained from a pool of hermaphroditic individuals, corresponding to the 1,8-cineole-rich chemotype. The peak area represents its relative abundance. Retention times (minutes) are represented in the x-axis. Only the 14 identified peaks are numbered (details are shown in **Table 1**). Radar charts displaying the mean relative abundance (0–100 scale) of the 14 identified compounds for 1,8-cineole rich **(B)** and linalool-rich (**C**) chemotypes. Each axis corresponds to one compound.

**Table 1.** Essential oils identified in *Thymus* sect. *Mastichina*. Compounds were identified based on their retention times (min) and characteristic mass fragments (m/z). Their classification according to chemical nature and relative abundance (mean percentage ± SD) are also provided. Values in parentheses indicate the range (minimum–maximum) calculated from population means. Compounds are listed in order of elution.

| Compound | Retention time (min) | m/z | id | Chemical nature | Relative abundance (%) |
| --- | --- | --- | --- | --- | --- |
| 1 | 5.37 | 136 | α-pinene | Monoterpene hydrocarbons | 2.577 ± 0.956<br>(1.175 – 4.873) |
| 2 | 5.75 | 136 | camphene | Monoterpene hydrocarbons | 2.048 ± 2.518<br>(0 – 9.883) |
| 3 | 6.28 | 136 | sabinene | Monoterpene hydrocarbons | 1.732 ± 0.703<br>(0.323 – 2.858) |
| 4 | 6.4 | 136 | β-pinene | Monoterpene hydrocarbons | 2.542 ± 0.778<br>(1.315 – 3.732) |
| 5 | 6.67 | 136 | β-myrcene | Monoterpene hydrocarbons | 0.733 ± 0.436<br>(0 – 1.864) |
| 6 | 7.7 | 136 | limonene | Monoterpene hydrocarbons | 0.969 ± 0.519<br>(0.397 – 2.691) |
| 7 | 7.79 | 154 | 1,8-cineole | Oxygenated monoterpenes | 71.13 ± 13.890<br>(35.594 – 89.487) |
| 8 | 8.81 | 154 | sabinene hydrate | Oxygenated monoterpenes | 1.150 ± 0.543<br>(0.651 – 3.505) |
| 9 | 9.61 | 154 | linalool | Oxygenated monoterpenes | 5.543 ± 10.40<br>(0 – 42.845) |
| 10 | 10.94 | 152 | camphor | Oxygenated monoterpenes | 2.819 ± 5.181<br>(0 – 20.287) |
| 11 | 11.59 | 154 | δ-terpineol | Oxygenated monoterpenes | 0.884 ± 0.333<br>(0 – 1.549) |
| 12 | 11.64 | 154 | borneol | Oxygenated monoterpenes | 3.868 ± 4.359<br>(0 – 13.261) |
| 13 | 12.29 | 154 | α-terpineol | Oxygenated monoterpenes | 3.456 ± 1.386<br>(1.086 – 5.849) |
| 14 | 13.62 | 196 | linalool acetate | Oxygenated monoterpenes | 0.543 ± 3.092<br>(0 – 18.504) |
| | | | | Monoterpene hydrocarbons | 89.40% ± 3.458<br>(77.67% – 95.90%) |
| | | | | Oxygenated monoterpenes | 10.60% ± 3.458<br>(4.096% – 22.33%) |

The most common chemotype was characterized by high levels of 1,8-cineole (also known as eucalyptol; compound 7), with a mean concentration of 71.13% ± 13.09 (**Figure 2A and B**; **Table 1)**. When 1,8-cineole reached or exceeded this average value, the remaining compounds individually rarely accounted for more than 5% to the total composition. While the average concentration of linalool (compound 9) was 5.54% ± 10.40 (**Table 1**), in seven populations it accounted for more than 10% of the total aromatic compounds (**Table S2**). In two of these populations, A01 and M09, both diploids and located in the southern Iberian Peninsula, the population average concentration of linalool reached the levels of 1,8-cineole, with 42.85% and 29.46% for linalool, respectively, compared to 41.59% and 35.55% for 1,8-cineole. Moreover, the hermaphroditic pool from population A01 exhibited a chemotype clearly dominated by linalool (72.0%) instead of 1,8-cineole (19.05%) (**Figure 2C; Table S2**). Linalool acetate (compound 14) was generally a minor constituent of the EOs (0.54% ± 3.09; Table 1), but in the M09 population it accounted for 18.50% of the total compounds. Other oxygenated monoterpenes, such as camphor and borneol (compounds 10 and 12) also exhibited marked interpopulation differences. Although the overall mean concentrations of these compounds were 2.82% ± 5.18 and 3.87% ± 4.36, respectively (**Table 1**), six populations from the southwestern Iberian Peninsula (A08, D04, D05, D07, D08, and M11, all from the ‘*Doñana’* diploid group) stood out for consistently accumulating >10% of one or both compounds (**Table S2**). Finally, camphene (compound 2), a minor monoterpene hydrocarbon (2.05% ± 2.52; **Table 1**), reached approximately 10% of the total aromatic profile in three populations from the ‘*Doñana’* group (D05, D07, and D08; **Table S2**).

Further qualitative analysis of 13 pools exhibiting the alternative linalool-enriched chemotype revealed considerable variability among individuals within populations. While in some populations only one individual produced linalool at high concentrations, in others, up to four out of five individuals accumulated substantial amounts of this compound (**Table S3**). Moreover, within some pools, we observed individuals exhibiting distinct chemotypes: some dominated by 1,8-cineole, others dominated by linalool, and yet others with a more balanced composition of both compounds. In certain cases, up to two or three different chemotypes coexisted within the same pool (**Table S3**).

### Variation in essential oil composition across ploidy levels and genetic groups

To analyze differences in chemical composition between diploid and tetraploid individuals, we first used the relative proportions of all 14 individual compounds. This initial approach revealed notable differences in essential oil composition between ploidy levels (MANOVA: Wilks’ λ = 0.362; df = 1; P < 0.001). Significant differences were observed in seven out of the 14 aromatic compounds. The relative abundances of α-pinene, camphene, sabinene hydrate, camphor, and borneol were higher in diploid individuals, whereas limonene and linalool acetate were more abundant in tetraploids. Linalool, by contrast, did not differ significantly between ploidy levels, whereas 1,8-cineole showed a trend toward significance (P = 0.076), with slightly higher concentrations in tetraploids (**Figure 3A**). When EOs were grouped based on their chemical nature, significant differences between ploidy levels were also found (MANOVA: Wilks’ λ = 0.902; df = 1; P = 0.021): monoterpene hydrocarbons were more abundant in diploids, while oxygenated monoterpenes accounted for a higher proportion of compounds in tetraploids (**Figure S1A**).

**Figure 3.**
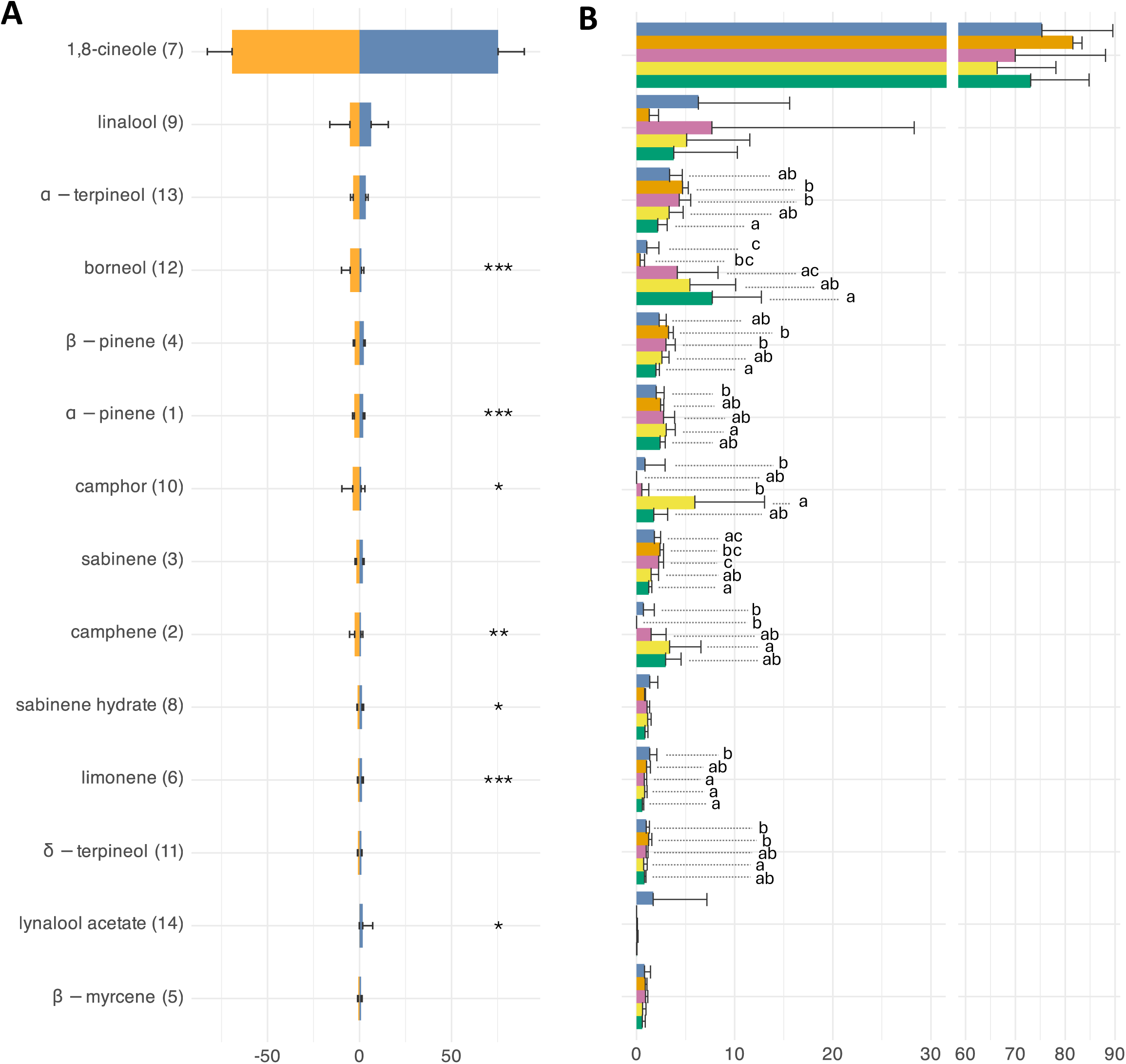
Horizontal bar plots depicting the relative abundances (mean ± SD) of the 14 identified EOs, arranged in descending order of mean abundance. Parentheses indicate the compound labels as shown in Figure 2A. Comparisons were made between **(A)** diploid (orange) and tetraploid (blue) individuals, and **(B)** the five genetic groups described in García-Cárdenas et al. (2025): ‘*Tetraploid’* (blue), ‘*Hercynian’* (orange), ‘*Algarve’* (green), ‘*Doñana’* (yellow), and ‘*Cadiz*’ (pink). * < 0.05, ** < 0.01, *** < 0.001. Different letters in (**B**) indicate significant differences at the 0.05 level.

We also detected differences in chemical composition among genetic groups, based on the relative proportions of all 14 individual aromatic compounds (MANOVA: Wilks’ λ = 0.121; df = 4; P < 0.001). Nine of them showed significant variation across genetic clusters: α-pinene, camphene, sabinene, b-pinene, limonene, camphor, d-terpineol, borneol, and α - terpineol. The ‘*Doñana’* populations exhibited notably higher relative abundances of α-pinene, camphene, and camphor compared to the ‘*Tetraploid’* group, with camphor in particular reaching much higher levels in ‘*Doñana’* than in any other genetic group.

Tetraploid populations, in contrast, were characterized by elevated levels of limonene, while δ-terpineol concentrations were most abundant in both the ‘*Hercynian’* and ‘*Tetraploid’* groups. Borneol levels were substantially high in southwestern populations, specifically in the ‘*Algarve’*, ‘*Cadiz’*, and ‘*Doñana’* groups. Additionally, sabinene, β-pinene, and α-terpineol showed reduced relative abundances in the ‘*Algarve’* populations but were comparatively higher in ‘*Cadiz’* and the ‘*Hercynian’* groups. Despite these compound-specific differences, no clear or consistent pattern of compound distribution was observed across all genetic clusters (**Figure 3B**). Finally, when compounds were grouped according to their chemical nature, we did not detect any significant differences among genetic clusters (MANOVA: Wilks’ λ = 0.830; df = 4; P = 0.089) (**Figure S1B**).

### Assessing climatic and edaphic effects on chemical profiles

Correlations between aromatic compounds and environmental variables revealed some statistically significant associations at the population level (**Figure 4A**). First, a notable pattern emerged with bio1 (annual mean temperature), which was positively correlated with the relative abundance of camphor, borneol, camphene, and α-pinene (*r_S_* ranging from 0.359 to 0.522, *P* < 0.021; **Table S4**), but negatively correlated with 1,8-cineole (*r_S_* = –0.40, *P* < 0.001). Second, the most relevant results were found for bio14 (precipitation of driest month) and bio17 (precipitation of driest quarter). Specifically, we observed significant negative correlations between these precipitation variables and the proportion of borneol, camphor, camphene, and α-pinene (*rS* ranging from –0.37 to –0.60, *P* < 0.010; **Table S4**), with the only exception of α-pinene, which showed a marginally significant correlation with bio17 (*P* = 0.052). Linalool exhibited a significant negative correlation with MRH (*r_S_*= –0.52, *P* < 0.001; **Figure 4A**). Overall, these patterns indicate that these compounds tend to accumulate under drier conditions. In contrast, 1,8-cineole showed a positive correlation with bio14 (*r_S_* = 0.45, *P* < 0.001), suggesting greater abundance in populations from areas with higher minimum precipitation. Among the soil variables, few consistent patterns emerged, but WRP (water retention potential) showed a negative correlation with borneol and camphene (*r_S_* = –0.45 and –0.44, respectively, *P* = 0.010), whereas linalool was positively correlated with organic matter (OM) (*r_S_* = 0.41, P = X; **Table S4**).

**Figure 4.**
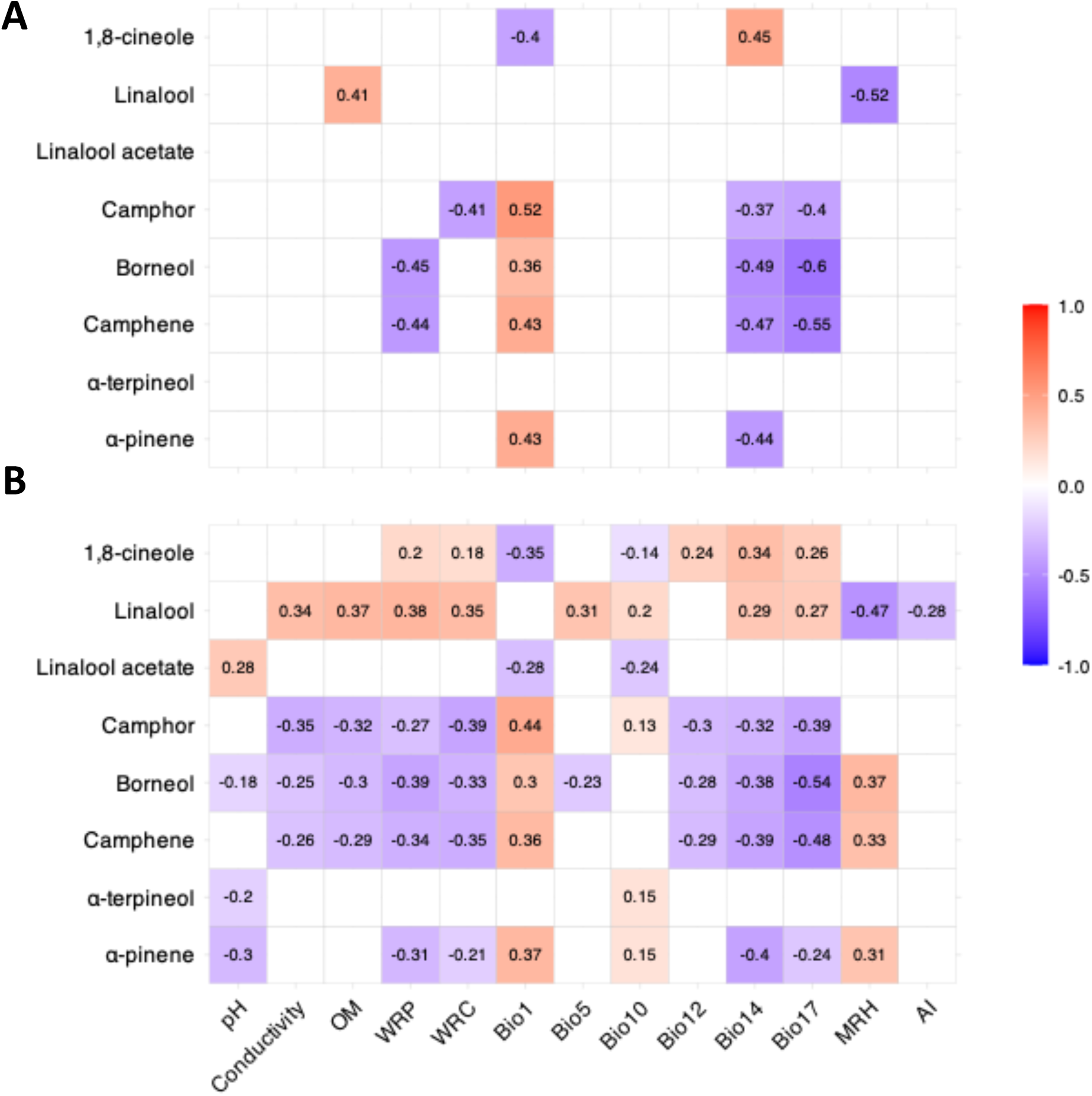
Correlograms showing significant correlations between selected aromatic compounds and environmental variables at the population (**A**) and individual (**B**) levels. Values indicate Spearman’s r for significant correlations after Bonferroni correction. Colors represent the strength and direction of correlations, following a gradient from –1 (blue) to +1 (red). The analyses included all soil-related variables (pH, conductivity, WRP, WRC, and OM) and eight climatic variables: bio1 (annual mean temperature), bio5 (maximum temperature of warmest month), bio10 (mean temperature of warmest quarter), bio12 (annual precipitation), bio14 (precipitation of driest month), bio17 (precipitation of driest quarter), MRH (mean relative humidity), and AI (aridity index).

At the individual level, these general patterns were largely maintained. Camphor, borneol, camphene, and α-pinene remained positively associated with temperature (bio1) but negatively associated with precipitation variables (bio12 – annual precipitation, bio14, and bio17). In contrast, 1,8-cineole displayed opposite trends, while linalool showed negative correlations with bio14 and bio17. Camphor, borneol and camphene were negatively correlated with almost all edaphic variables (conductivity, OM, WRP - water retention potential, and WRC - water retention capacity), while linalool showed positive correlations with OM, WRP, and WRC (**Figure 4B; Table S5**).

## Discussion

In this study, we provide an integrative overview of EO profiles in *Thymus* sect. *Mastichina*, identifying key patterns of chemical diversity and their potential drivers. First, we detected moderate chemical diversity both within and among populations: although many populations exhibited 1,8-cineol-enriched profiles, others were dominated by linalool, and in some cases multiple chemotypes coexisted within the same population. Second, considerable variation was observed among populations, with moderate differences between diploid and tetraploid individuals and among genetically defined clusters, pointing to potential effects of ploidy and genetic background on EO composition. While the major compounds—1,8-cineole and linalool—remained relatively stable across all populations, other minor constituents showed considerable variation across populations. Third, environmental factors also contributed to shaping EO profiles, suggesting that local climatic and edaphic conditions modulate terpene biosynthesis. Together, these findings suggest a complex interplay of genetic, ploidy, and environmental influences in determining chemotypic diversity in *Thymus* sect. *Mastichina*, providing a robust framework for understanding the ecological and evolutionary drivers of EO variation in this group.

The chemical composition of EOs in *Thymus* sect. *Mastichina* displays both conserved and divergent patterns across its geographic range. Although the MANOVA results revealed significant differences in chemical composition among populations, the two most abundant compounds — 1,8-cineole and linalool — were consistently present across all populations and did not vary significantly with ploidy level or genetic groups. This consistency in compound presence does not preclude variation in their relative abundance across populations or even among individuals within the same stand. While most populations are composed of individuals sharing a single chemotype, others have two or more chemotypes, highlighting a high level of inter- and intraspecific chemical variability that is consistent with the heterogeneous distribution of chemotypes reported in other *Thymus* species (e.g., Adzet *et al*. 1977; Thompson *et al*. 2003; Keefover-Ring *et al*. 2009). Additionally, there is a considerable increase of the relative abundance of linalool in seven populations from the southwestern Iberian Peninsula, along with increased levels of typically minor constituents such as borneol and camphor. Yet, a notable exception is the ‘*Doñana’* genetic group, in which borneol concentrations are consistently higher, as reported in previous studies (e.g., Salgueiro *et al*. 1997; Figueiredo *et al*. 2008). This indicates a relatively stable chemotypic profile and suggests that this lineage may retain characteristic chemical signatures, likely reflecting a combination of phylogenetic distinctiveness and ecological factors (Baser & Buchbauer 2009; Etri & Pluhár 2024; Ouarghidi *et al*. 2025).

The synthesis of individual monoterpenes is unlikely to occur independently. In *T. vulgaris*, Thompson et al. (2003) showed that the production of one compound can inversely affect the levels of others, reflecting an interdependent metabolic network. This trade-off likely contributes to the negative correlation observed between 1,8-cineole and linalool (Kampranis *et al*. 2007). The relative flux through these competing branches can be modulated by the activity of specific monoterpene synthases (Bataillon *et al*. 2022), which in turn may be influenced by allelic variation, gene expression or transcriptional regulation (Thompson *et al*. 2003; Rabiei *et al*. 2018; Bao *et al*. 2023). In particular, the strong association between a small number of terpene-synthase loci and major chemotypic differences observed in *T. vulgaris* (Bataillon *et al*. 2022) suggests that limited genetic architectures can underlie diverse chemical phenotypes. Such enzymatic and genetic regulation may provide the basis for the observed intraspecific chemical variability in *Thymus* sect. *Mastichina*, including the coexistence of multiple chemotypes within the same population (Benabdelkader *et al*. 2015).

Environmental conditions, alongside genetic factors, have been shown to modulate monoterpene biosynthesis in *Thymus* species (Salgueiro *et al*. 1995, 1997; Miguel *et al*. 2004; Amiot *et al*. 2005; Thompson *et al*. 2007). In *T. vulgaris*, chemotypes exhibit a well-defined spatial distribution, putatively maintained by local ecological pressures such as herbivory, drought, and extreme temperatures (Thompson *et al*. 1998, 2003, 2007).

Similarly, populations of *T. carnosus*, *T. mastichina*, and *T. albicans* along the Atlantic coast, exposed to maritime humidity, tend to accumulate higher levels of linalool or borneol (Salgueiro *et al*. 1995, 1997). In our study, correlation between aromatic profiles and climatic variables indicated that camphene and its derivatives, borneol and camphor, were more prominent in populations from drier sites, suggesting shifts in secondary metabolite allocation under water-limited conditions (Ramezani *et al*. 2020), whereas 1,8-cineole was generally more abundant in wetter environments. While studies in *Salvia* support a down-regulation of cineole synthase, the enzyme directly responsible for 1,8-cineole biosynthesis, under drought stress (Radwan *et al*. 2017), other reports show an opposite pattern, with 1,8-cineole concentrations increasing during the drought period in Eastern Iberian populations of *T. vulgaris* (Llorens-Molina & Vacas 2017), indicating that the response of this compound to water stress may be species- or context-specific.

The patterns observed in this study indicate that variation in EO composition may not be random, but could reflect adaptive responses to local environmental conditions, mediated by both metabolic regulation and ecological pressures. The functional roles of individual compounds support this hypothesis: 1,8-cineole is primarily associated with antimicrobial, antifungal, antioxidant, or insecticide activity, while linalool plays a broader role, contributing to pollinator attraction via floral scent while also offering antifungal and insect-repellent functions (Dorman *et al*. 2000; Faleiro *et al*. 2003; Pina-Vaz *et al*. 2004; Figueiredo *et al*. 2008; Azerad 2014; Bajalan *et al*. 2017; Rodrigues *et al*. 2020; Cai *et al*.

2021). Borneol and α-terpineol are particularly valued for its antibacterial properties, and camphor plays roles in oxidative stress protection, antipathogen properties and herbivore deterrence (Bajalan *et al*. 2017). Taken together, our results suggest that the chemical diversity observed in *Thymus* sect. *Mastichina* likely arises from the interplay between environmental pressures and intrinsic metabolic pathways, reflecting adaptive strategies that optimize ecological interactions and population resilience under variable climatic conditions. Nonetheless, the potential role of phenotypic plasticity in generating this chemical diversity cannot be ruled out and should be explicitly tested in future studies.

## Conclusions

Our results provide a comprehensive view of chemotypic diversity and ecological adaptation in *Thymus* sect. *Mastichina*, even though their value for species delimitation remains limited. Although differences between diploid and tetraploid individuals were moderate, ongoing tetraploid introgression into diploid populations may further blur these boundaries, reinforcing the chemical continuity observed across taxa. Moreover, the broad ecological functions of the major monoterpenes indicate that the major variation documented in this study is unlikely to be arbitrary; instead, it likely reflects adaptive strategies that enhance plant performance and modulate ecological interactions across contrasting environmental conditions. The coexistence of multiple chemotypes within populations, together with overlapping chemical signatures across taxa, highlights the evolutionary complexity of the group and emphasizes the need to combine chemical, genetic, and ecological evidence to achieve a robust taxonomic resolution.

## Acknowledges

The authors thank to Francisco Javier Jiménez-López, Javier López-Tirado, Laura Serrano, Laura Fernández, Andrés Melero and Billy Williams-Marland for their field assistant. We particularly thank the anonymous referees for their helpful comments. This study is part of the Conserva3 project (TED2021-130133B-I00) funded by MCIN/AEI/10.13039/501100011033 to R.B.P. and D.N.L. with funds from the European Union “NextGenerationEU”/PRTR.

## Author contributions

RBP and DNL conceived the study and established the methodology. JCV, EMC, DDP, MOH, DNL and RBP carried out the sampling. DDP, MOH, DNL obtained and analyzed the soil and climatic data. JCV, EMC, HP, PG and BE obtained and analyzed the biochemical data. JCV, EMC, DNL and RBP performed the statistical analyses. JCV, DNL and RBP drafted the manuscript. All authors contributed to the final manuscript and approved the submitted version.

