## Supplementary tables and figures for "Ecological and Genetic Determinants of Essential Oil Diversity in Mediterranean *Thymus*"

[1] Departamento de Biología Vegetal y Ecología, Universidad de Sevilla, Sevilla, Spain.

[2] CEFE, Univ. Montpellier, CNRS, EPHE, IRD, Montpellier, France.

[3] Institut for Ecoscience Terrestrisk Økologi, Århus, Denmark.

[4] Departamento de Botánica, Ecología y Fisiología Vegetal, Universidad de Córdoba, Córdoba, Spain.

### Supplementary material

**Figure S1.** Violin plots combined with boxplots show the full distribution of values obtained for the relative frequencies on the EOs based on their chemical nature. Comparisons were made **(A)** between diploid and tetraploid individuals **(B)**, and among the five genetic groups described in García-Cárdenas et al. (2025): '*Tetraploid'* (blue), '*Hercynian'* (orange), '*Algarve'* (green), '*Doñana'* (yellow), and '*Cadiz*' (pink). Boxplots represent the interquartile range (IQR); the thin black line indicates 1.5× the IQR, the central line shows the median, and the black dot represent the mean sample value; black dots outside the IQR represent the outlayer values. **, $P < 0.010$.

**Table S1**. Populations of *Thymus* sect. *Mastichina*, indicating their genetic cluster (*'Tetraploid'*, *'Hercynian', 'Algarve'*, *'Doñana'*, and *'Cadiz'*), ploidy level (estimated through flow cytometry; García-Cárdenas et al., in rev.), and collection site. Populations are ordered alphabetically.

| Population | Genetic cluster | Ploidy | Locality | Coordinates | Herbarium voucher (SEV) |
|---|---|---|---|---|---|
| **A01** | Cadiz | 2x | Spain. Cádiz, Puerto Real. Dehesa de las Yeguas | 36.5557, -6.1351 | SEV290208 |
| **A02** | Cadiz | 2x | Spain. Cádiz, la Barrosa Periurban Park | 36.3628, -6.1704 | SEV290209 |
| **A03** | Cadiz | 2x | Spain. Cádiz, Conil de la Frontera. ZEC Pinar de Roche | 36.3163, -6.1333 | SEV290210 |
| **A04** | Doñana | 2x | Spain. Sevilla. Villamanrique de la Condesa. Doñana Natural Park. La Juncosilla | 37.1916, -6.3294 | SEV290211 |
| **A05** | Algarve | 2x | Portugal. Algarve, Faro. Ria Formosa Natural Park | 37.0288, -7.9773 | SEV290212 |
| **A06** | Doñana | 2x | Spain. Huelva, Villarrasa. Dehesa del Duque | 37.3563, -6.6044 | SEV290213 |
| **A07** | Doñana | 2x | Spain. Huelva, Bonares | 37.2586, -6.7254 | SEV290214 |
| **A08** | Doñana | 2x | Spain. Doñana Natural Park. Pinar de la Algaida | 36.8553, -6.3075 | SEV290215 |
| **A09** | Algarve | 2x | Portugal. Algarve, Quarteira. Vale do Lobo | 37.0617, -8.0796 | SEV290216 |
| **A10** | Algarve | 2x | Portugal. Algarve, Almancil. Vale do Garrao. Ria Formosa Natural Park | 37.0443, -8.0382 | SEV290217 |
| **A11** | Doñana | 2x | Spain. Huelva, Niebla. Rabo Conejo | 37,4887, -6,7192 | SEV290218 |
| **A12** | Cadiz | 2x | Spain. Cádiz, Vejer de la Frontera. Las Lomas | 36.2988, -5.9087 | SEV290219 |
| **A13** | Cadiz | 2x | Spain. Cádiz, Vejer de la Frontera. El Soto | 36.2422, -5.9287 | SEV290220 |
| **A14** | Cadiz | 2x | Spain. Cádiz, Chiclana. Pinar del Hierro | 36.3880, -6.1222 | SEV290221 |

| | | | | | |
|---|---|---|---|---|---|
| **A15** | Doñana | 2x | Spain. Sevilla. Villamanrique de la Condesa. Doñana Natural Park. Dehesa de Gato | 37.2540, -6.3395 | SEV290222 |
| **D01** | Doñana | 2x | Spain. Huelva, Almonte. El Porretal. ZEC Doñana North and West | 37.1671, -6.4967 | SEV290223 |
| **D02** | Doñana | 2x | Spain. Huelva, Hinojos. ZEC Doñana North and West | 37.2865, -6.4453 | SEV290224 |
| **D04** | Algarve | 2x | Portugal. Algarve, Quelfes. Pinheiros do Marim. Ria Formosa Natural Park | 37.0336, -7.8181 | SEV290226 |
| **D05** | Doñana | 2x | Spain. Huelva, Abalario. Doñana Natural Park | 37.0719, -6.6506 | SEV290227 |
| **D06** | Doñana | 2x | Spain. Huelva, Ribetehilos. Doñana Natural Park | 37.1500, -6.6433 | SEV290228 |
| **D07** | Doñana | 2x | Spain. Huelva, Corral de la Liebre. Doñana National Park | 36.9079, -6.4141 | SEV290229 |
| **D08** | Doñana | 2x | Spain. Huelva. Corralillo Oscuro. Doñana National Park | 36.9869, -6.5215 | SEV290230 |
| **M01** | Tetraploid | 4x | Spain. Sevilla, Cazalla de la Sierra. Sierra Morena de Sevilla Natural Park | 37.9075, -5.8023 | SEV290231 |
| **M02** | Hercynian | 2x | Spain. Córdoba, El Vacar. ZEC Guadalmellato | 38.1001, -4.8330 | SEV290232 |
| **M03** | Doñana | 2x | Spain. Huelva, Cartaya. Campo Común de Abajo | 37.2383, -7.0796 | SEV290233 |
| **M04** | Tetraploid | 4x | Spain. Sevilla, Gilena | 37.2805, -4.9014 | SEV290234 |
| **M05** | Tetraploid | 4x | Spain. Sevilla, Gerena | 37.5611, -6.1593 | SEV290235 |
| **M06** | Tetraploid | 4x | Portugal. Algarve, Monte Gordo | 37.1855, -7.4880 | SEV290236 |
| **M07** | Tetraploid | 4x | Spain. Madrid. Hoyos de Manzanares. Cuenca Alta del Manzanares Regional Park | 40.6021, -3.9167 | SEV290237 |
| **M08** | Tetraploid | 4x | Spain. Cádiz, Villaluenga del Rosario. Sierra de Grazalema Natural Park | 36.8108, -5.3907 | SEV290238 |
| **M09** | Tetraploid | 4x | Spain. Málaga, Ronda, Pinsapar de la Nava. Sierra de las Nieves Natural Park | 36.6650, -5.0510 | SEV290239 |
| **M10** | Doñana | 2x | Spain. Huelva, Cartaya. Campo Común de Arriba | 37.3299, -7.0997 | SEV290240 |
| **M11** | Doñana | 2x | Spain. Huelva, El Portil | 37.2035, -7.0136 | SEV290241 |
| **M12** | Tetraploid | 4x | Spain. Cuenca, Pajaroncillo | 39.9301, -1.7257 | SEV290242 |
| **M13** | Tetraploid | 4x | Spain. Cuenca, San Clemente | 39.3376, -2.4718 | SEV290243 |
| **M14** | Hercynian | 2x | Portugal. Guarda, Serra da Estrela Natural Park | 40.3788, -7.5066 | SEV290244 |
| **M15** | Tetraploid | 4x | Spain. León, Ponferrada | 42.5639, -6.7705 | SEV290245 |
| **M16** | Tetraploid | 4x | Spain. Zamora, Toro | 41.5160, -5.4148 | SEV290246 |
| **M18** | Tetraploid | 4x | Spain. Albacete, Parideras | 38.5817, -2.5262 | SEV290492 |

**Table S2.** Mean relative concentrations (%) of essential oils identified in *Thymus* sect. *Mastichina* at the population level. Values are shown for the population mean ("Mean"), female (F) and hermaphroditic (H) pools. Ploidy level is indicated in parentheses. As data were obtained from pooled samples, standard deviation values are not provided. Compounds are listed in order of elution (see **Table 1** for compound identification and related information).

| **Pop** | **Group** | **α-pinene** | **Camphene** | **Sabinene** | **β-pinene** | **β-myrcene** | **Limonene** | **1,8-cineole** | **Sabinene hydrate** | **Linalool** | **Camphor** | **δ-terpineol** | **Borneol** | **α-terpineol** | **Linalool acetate** |
|---|---|---|---|---|---|---|---|---|---|---|---|---|---|---|---|
| A01 (2x) | Mean | 1.35 | 1.16 | 1.52 | 1.31 | 0.47 | 0.40 | 41.59 | 1.34 | 42.85 | 0.75 | 0.69 | 4.28 | 2.17 | 0.14 |
| | F | 1.77 | 1.89 | 1.98 | 1.68 | 0.52 | 0.49 | 64.13 | 1.57 | 13.69 | 1.49 | 0.88 | 7.23 | 2.68 | 0.00 |
| | H | 0.93 | 0.43 | 1.05 | 0.95 | 0.42 | 0.31 | 19.05 | 1.11 | 72.00 | 0.00 | 0.49 | 1.33 | 1.66 | 0.27 |
| A02 (2x) | Mean | 3.75 | 3.15 | 2.66 | 3.38 | 0.96 | 0.94 | 65.93 | 1.28 | 1.86 | 0.61 | 1.14 | 9.60 | 4.76 | 0.00 |
| | F | 4.02 | 3.68 | 2.58 | 3.40 | 0.87 | 0.93 | 64.36 | 1.25 | 1.90 | 0.91 | 1.21 | 10.38 | 4.52 | 0.00 |
| | H | 3.48 | 2.62 | 2.74 | 3.36 | 1.05 | 0.95 | 67.50 | 1.31 | 1.82 | 0.30 | 1.07 | 8.82 | 5.00 | 0.00 |
| A03 (2x) | Mean | 4.26 | 3.66 | 2.56 | 3.73 | 1.04 | 1.11 | 65.79 | 1.29 | 0.48 | 1.68 | 1.13 | 8.59 | 4.68 | 0.00 |
| | F | 4.16 | 3.58 | 2.56 | 3.70 | 1.04 | 1.06 | 66.53 | 1.27 | 0.48 | 1.96 | 1.17 | 7.89 | 4.61 | 0.00 |
| | H | 4.37 | 3.73 | 2.56 | 3.76 | 1.05 | 1.16 | 65.06 | 1.30 | 0.47 | 1.40 | 1.10 | 9.29 | 4.75 | 0.00 |
| A04 (2x) | Mean | 2.89 | 0.75 | 2.42 | 3.44 | 1.00 | 1.26 | 67.24 | 1.23 | 11.64 | 0.80 | 1.09 | 1.04 | 4.96 | 0.24 |
| | F | 2.98 | 0.79 | 2.49 | 3.46 | 1.09 | 1.06 | 65.98 | 1.22 | 12.78 | 0.74 | 1.01 | 1.15 | 4.75 | 0.49 |
| | H | 2.80 | 0.70 | 2.34 | 3.42 | 0.92 | 1.47 | 68.49 | 1.24 | 10.51 | 0.85 | 1.17 | 0.93 | 5.16 | 0.00 |
| A05 (2x) | Mean | 2.19 | 1.49 | 1.37 | 2.31 | 0.71 | 0.45 | 83.69 | 0.88 | 0.00 | 0.30 | 0.68 | 3.51 | 2.42 | 0.00 |
| | F | 1.83 | 1.71 | 1.11 | 1.97 | 0.57 | 0.36 | 84.20 | 0.74 | 0.00 | 0.61 | 0.67 | 4.10 | 2.13 | 0.00 |
| | H | 2.55 | 1.27 | 1.63 | 2.65 | 0.86 | 0.54 | 83.18 | 1.02 | 0.00 | 0.00 | 0.68 | 2.93 | 2.72 | 0.00 |
| A06 (2x) | Mean | 2.32 | 2.22 | 0.98 | 2.15 | 0.41 | 0.64 | 75.21 | 1.47 | 4.69 | 2.71 | 1.03 | 3.90 | 2.28 | 0.00 |
| | F | 3.12 | 2.16 | 1.96 | 3.05 | 0.83 | 0.77 | 77.23 | 1.01 | 0.00 | 2.83 | 1.04 | 2.61 | 3.39 | 0.00 |
| | H | 1.51 | 2.27 | 0.00 | 1.26 | 0.00 | 0.51 | 73.18 | 1.93 | 9.38 | 2.59 | 1.02 | 5.18 | 1.17 | 0.00 |

| | | | | | | | | | | | | | | | |
|---|---|---|---|---|---|---|---|---|---|---|---|---|---|---|---|
| A07 (2x) | Mean | 2.58 | 1.99 | 1.14 | 2.34 | 0.37 | 0.60 | 81.52 | 1.01 | 0.00 | 3.34 | 0.67 | 1.98 | 2.47 | 0.00 |
| | F | 2.13 | 1.75 | 0.79 | 1.88 | 0.00 | 0.44 | 84.45 | 0.89 | 0.00 | 3.33 | 0.60 | 1.78 | 1.95 | 0.00 |
| | H | 3.04 | 2.23 | 1.48 | 2.80 | 0.75 | 0.76 | 78.58 | 1.12 | 0.00 | 3.36 | 0.74 | 2.17 | 2.98 | 0.00 |
| A08 (2x) | Mean | 2.88 | 4.04 | 1.05 | 2.02 | 0.54 | 0.99 | 60.53 | 0.78 | 3.65 | 10.79 | 0.33 | 9.15 | 3.27 | 0.00 |
| | F | 2.92 | 3.43 | 1.19 | 2.31 | 0.65 | 1.16 | 67.66 | 0.61 | 1.30 | 8.95 | 0.28 | 6.03 | 3.50 | 0.00 |
| | H | 2.84 | 4.64 | 0.91 | 1.73 | 0.42 | 0.82 | 53.40 | 0.95 | 5.99 | 12.62 | 0.38 | 12.27 | 3.03 | 0.00 |
| A09 (2x) | Mean | 2.77 | 3.83 | 1.44 | 1.88 | 0.83 | 0.68 | 58.36 | 1.12 | 13.86 | 3.32 | 0.82 | 9.04 | 2.06 | 0.00 |
| | F | 3.03 | 4.86 | 1.20 | 1.78 | 0.81 | 0.66 | 50.26 | 1.45 | 17.16 | 1.93 | 0.90 | 14.20 | 1.77 | 0.00 |
| | H | 2.52 | 2.81 | 1.68 | 1.97 | 0.84 | 0.70 | 66.46 | 0.79 | 10.56 | 4.70 | 0.75 | 3.87 | 2.35 | 0.00 |
| A10 (2x) | Mean | 2.04 | 1.87 | 1.30 | 1.95 | 0.60 | 0.63 | 78.48 | 0.69 | 1.29 | 1.57 | 0.87 | 5.73 | 2.93 | 0.05 |
| | F | 1.48 | 1.37 | 1.15 | 1.56 | 0.41 | 0.42 | 85.37 | 0.46 | 1.78 | 1.18 | 0.69 | 2.35 | 1.77 | 0.00 |
| | H | 2.60 | 2.36 | 1.46 | 2.34 | 0.79 | 0.84 | 71.59 | 0.91 | 0.80 | 1.97 | 1.04 | 9.11 | 4.08 | 0.10 |
| A11 (2x) | Mean | 2.31 | 0.00 | 1.91 | 2.95 | 0.86 | 0.70 | 67.62 | 1.02 | 17.48 | 0.00 | 0.86 | 0.04 | 4.01 | 0.25 |
| | F | 2.13 | 0.00 | 1.77 | 2.74 | 0.79 | 0.73 | 67.71 | 0.98 | 18.51 | 0.00 | 0.79 | 0.08 | 3.68 | 0.10 |
| | H | 2.49 | 0.00 | 2.05 | 3.15 | 0.94 | 0.66 | 67.53 | 1.07 | 16.45 | 0.00 | 0.92 | 0.00 | 4.35 | 0.39 |
| A12 (2x) | Mean | 2.27 | 0.00 | 2.34 | 3.32 | 1.08 | 0.75 | 82.63 | 0.88 | 0.19 | 0.00 | 1.06 | 0.11 | 5.37 | 0.00 |
| | F | 2.90 | 0.00 | 2.81 | 4.03 | 1.25 | 0.85 | 80.23 | 0.92 | 0.21 | 0.00 | 1.10 | 0.10 | 5.60 | 0.00 |
| | H | 1.64 | 0.00 | 1.86 | 2.60 | 0.91 | 0.64 | 85.03 | 0.85 | 0.18 | 0.00 | 1.02 | 0.12 | 5.14 | 0.00 |
| A13 (2x) | Mean | 1.88 | 0.45 | 2.05 | 2.73 | 0.98 | 0.91 | 82.15 | 0.97 | 0.28 | 0.23 | 1.05 | 1.14 | 5.20 | 0.00 |
| | F | 2.21 | 0.71 | 2.19 | 2.97 | 1.06 | 0.83 | 80.32 | 0.97 | 0.33 | 0.31 | 1.10 | 1.61 | 5.40 | 0.00 |
| | H | 1.54 | 0.18 | 1.91 | 2.49 | 0.91 | 0.99 | 83.98 | 0.97 | 0.22 | 0.15 | 0.99 | 0.67 | 5.00 | 0.00 |
| A14 (2x) | Mean | 2.97 | 0.46 | 2.42 | 3.56 | 0.95 | 0.66 | 81.76 | 0.72 | 0.41 | 0.00 | 0.82 | 1.26 | 4.00 | 0.00 |
| | F | 3.02 | 0.92 | 2.27 | 3.27 | 0.89 | 0.61 | 81.05 | 0.73 | 0.41 | 0.00 | 0.83 | 2.26 | 3.75 | 0.00 |

| | | | | | | | | | | | | | | | |
|---|---|---|---|---|---|---|---|---|---|---|---|---|---|---|---|
| | H | 2.93 | 0.00 | 2.57 | 3.85 | 1.01 | 0.71 | 82.48 | 0.72 | 0.41 | 0.00 | 0.81 | 0.26 | 4.24 | 0.00 |
| A15 (2x) | Mean | - | - | - | - | - | - | - | - | - | - | - | - | - | - |
| | F | - | - | - | - | - | - | - | - | - | - | - | - | - | - |
| | H | - | - | - | - | - | - | - | - | - | - | - | - | - | - |
| D01 (2x) | Mean | 2.76 | 0.51 | 2.55 | 3.47 | 0.79 | 0.94 | 77.46 | 1.12 | 1.67 | 0.83 | 1.31 | 0.76 | 5.82 | 0.00 |
| | F | 2.81 | 0.40 | 2.57 | 3.58 | 0.85 | 1.21 | 76.80 | 1.11 | 1.49 | 0.52 | 1.43 | 0.69 | 6.53 | 0.00 |
| | H | 2.71 | 0.62 | 2.53 | 3.37 | 0.73 | 0.67 | 78.12 | 1.12 | 1.85 | 1.13 | 1.20 | 0.84 | 5.12 | 0.00 |
| D02 (2x) | Mean | 2.85 | 1.73 | 1.39 | 2.64 | 0.41 | 1.09 | 60.61 | 1.63 | 19.58 | 2.65 | 0.6545 | 1.75 | 3.04 | 0.00 |
| | F | 2.90 | 1.28 | 1.84 | 3.05 | 0.81 | 1.31 | 60.25 | 1.66 | 19.37 | 1.83 | 0.91 | 1.00 | 3.79 | 0.00 |
| | H | 2.79 | 2.19 | 0.94 | 2.23 | 0.00 | 0.87 | 60.97 | 1.60 | 19.79 | 3.46 | 0.38 | 2.49 | 2.30 | 0.00 |
| D04 (2x) | Mean | 2.60 | 4.71 | 0.88 | 1.79 | 0.26 | 0.61 | 71.75 | 0.77 | 0.00 | 1.87 | 0.87 | 12.61 | 1.29 | 0.00 |
| | F | 2.89 | 4.94 | 0.97 | 1.94 | 0.51 | 0.61 | 68.94 | 0.87 | 0.00 | 1.41 | 1.03 | 14.09 | 1.78 | 0.00 |
| | H | 2.32 | 4.47 | 0.78 | 1.64 | 0.00 | 0.61 | 74.56 | 0.67 | 0.00 | 2.32 | 0.72 | 11.12 | 0.80 | 0.00 |
| D05 (2x) | Mean | 4.87 | 9.88 | 0.93 | 2.44 | 0.27 | 1.04 | 46.93 | 1.23 | 0.45 | 16.85 | 0.00 | 13.26 | 1.83 | 0.00 |
| | F | 4.00 | 9.03 | 0.76 | 2.08 | 0.00 | 0.67 | 52.40 | 1.07 | 0.00 | 15.76 | 0.00 | 12.88 | 1.35 | 0.00 |
| | H | 5.75 | 10.74 | 1.10 | 2.80 | 0.54 | 1.41 | 41.47 | 1.39 | 0.91 | 17.94 | 0.00 | 13.65 | 2.32 | 0.00 |
| D06 (2x) | Mean | 3.63 | 2.30 | 2.38 | 3.61 | 1.08 | 0.95 | 73.82 | 1.12 | 0.15 | 3.00 | 0.89 | 2.73 | 4.33 | 0.00 |
| | F | 4.05 | 2.98 | 2.39 | 3.56 | 1.15 | 0.99 | 71.06 | 1.16 | 0.31 | 4.76 | 0.72 | 2.65 | 4.23 | 0.00 |
| | H | 3.20 | 1.61 | 2.38 | 3.66 | 1.01 | 0.91 | 76.58 | 1.09 | 0.00 | 1.24 | 1.06 | 2.82 | 4.43 | 0.00 |
| D07 (2x) | Mean | 3.46 | 8.74 | 0.32 | 1.58 | 0.23 | 0.63 | 45.61 | 1.38 | 5.33 | 20.29 | 0.61 | 10.75 | 1.09 | 0.00 |
| | F | 3.03 | 8.64 | 0.00 | 1.40 | 0.00 | 0.60 | 53.01 | 0.57 | 0.00 | 22.39 | 0.27 | 9.24 | 0.84 | 0.00 |
| | H | 3.88 | 8.84 | 0.65 | 1.76 | 0.45 | 0.65 | 38.21 | 2.19 | 10.66 | 18.18 | 0.95 | 12.25 | 1.33 | 0.00 |
| D08 (2x) | Mean | 4.18 | 8.70 | 0.82 | 2.15 | 0.47 | 0.70 | 50.00 | 1.13 | 2.05 | 19.36 | 0.58 | 7.98 | 1.87 | 0.00 |

| | | | | | | | | | | | | | | | |
|---|---|---|---|---|---|---|---|---|---|---|---|---|---|---|---|
| | F | 4.60 | 8.84 | 0.83 | 2.38 | 0.52 | 0.73 | 49.44 | 1.07 | 0.93 | 20.20 | 0.72 | 7.52 | 2.21 | 0.00 |
| | H | 3.76 | 8.56 | 0.81 | 1.93 | 0.43 | 0.66 | 50.55 | 1.19 | 3.17 | 18.52 | 0.44 | 8.45 | 1.54 | 0.00 |
| M01 (4x) | Mean | 1.80 | 0.82 | 1.61 | 1.92 | 0.79 | 0.55 | 83.10 | 0.97 | 2.58 | 0.42 | 1.15 | 1.63 | 2.67 | 0.00 |
| | F | 2.27 | 1.13 | 1.89 | 2.33 | 0.95 | 0.56 | 80.56 | 1.06 | 2.34 | 0.35 | 1.44 | 2.10 | 3.00 | 0.00 |
| | H | 1.34 | 0.50 | 1.32 | 1.51 | 0.63 | 0.53 | 85.63 | 0.87 | 2.81 | 0.49 | 0.87 | 1.16 | 2.33 | 0.00 |
| M02 (2x) | Mean | 2.21 | 0.00 | 2.27 | 2.91 | 0.84 | 1.33 | 80.49 | 0.85 | 2.12 | 0.00 | 1.52 | 0.75 | 4.73 | 0.00 |
| | F | 2.40 | 0.00 | 2.65 | 3.20 | 1.02 | 1.12 | 78.98 | 0.95 | 2.04 | 0.00 | 1.45 | 0.74 | 5.43 | 0.00 |
| | H | 2.02 | 0.00 | 1.88 | 2.62 | 0.65 | 1.54 | 81.99 | 0.74 | 2.20 | 0.00 | 1.58 | 0.76 | 4.03 | 0.00 |
| M03 (2x) | Mean | 2.22 | 2.00 | 1.93 | 2.32 | 0.78 | 0.76 | 72.77 | 0.97 | 2.20 | 0.97 | 0.53 | 8.22 | 4.26 | 0.09 |
| | F | 2.68 | 2.12 | 2.31 | 2.82 | 0.94 | 0.89 | 69.58 | 1.06 | 2.25 | 1.00 | 0.70 | 8.31 | 5.17 | 0.18 |
| | H | 1.75 | 1.87 | 1.54 | 1.81 | 0.62 | 0.63 | 75.96 | 0.87 | 2.15 | 0.94 | 0.36 | 8.14 | 3.35 | 0.00 |
| M04 (4x) | Mean | 2.28 | 0.00 | 2.48 | 3.08 | 0.83 | 1.10 | 80.32 | 1.19 | 2.19 | 0.00 | 1.41 | 0.00 | 5.13 | 0.00 |
| | F | 2.26 | 0.00 | 2.30 | 2.91 | 0.62 | 1.04 | 81.14 | 1.19 | 2.23 | 0.00 | 1.48 | 0.00 | 4.82 | 0.00 |
| | H | 2.30 | 0.00 | 2.66 | 3.24 | 1.04 | 1.15 | 79.50 | 1.18 | 2.15 | 0.00 | 1.35 | 0.00 | 5.43 | 0.00 |
| M05 (4x) | Mean | 2.59 | 0.00 | 2.86 | 3.62 | 0.99 | 0.85 | 78.57 | 1.19 | 1.94 | 0.00 | 1.55 | 0.00 | 5.85 | 0.00 |
| | F | 2.60 | 0.00 | 2.92 | 3.74 | 1.10 | 0.78 | 77.91 | 1.25 | 1.67 | 0.00 | 1.67 | 0.00 | 6.35 | 0.00 |
| | H | 2.57 | 0.00 | 2.79 | 3.49 | 0.89 | 0.93 | 79.23 | 1.13 | 2.21 | 0.00 | 1.42 | 0.00 | 5.34 | 0.00 |
| M06 (4x) | Mean | 3.52 | 3.19 | 2.54 | 2.83 | 0.74 | 1.95 | 67.17 | 2.24 | 2.57 | 6.56 | 1.11 | 2.07 | 3.51 | 0.00 |
| | F | 3.71 | 2.72 | 2.52 | 2.99 | 0.78 | 2.51 | 69.01 | 1.83 | 2.39 | 5.18 | 1.12 | 1.62 | 3.63 | 0.00 |
| | H | 3.33 | 3.67 | 2.55 | 2.67 | 0.71 | 1.39 | 65.33 | 2.66 | 2.74 | 7.94 | 1.11 | 2.53 | 3.38 | 0.00 |
| M07 (4x) | Mean | 1.57 | 0.65 | 2.08 | 1.87 | 0.00 | 1.03 | 80.52 | 1.12 | 6.85 | 0.00 | 0.82 | 0.98 | 2.18 | 0.33 |
| | F | 1.31 | 1.30 | 1.91 | 1.55 | 0.00 | 0.50 | 78.80 | 1.08 | 8.08 | 0.00 | 0.99 | 1.95 | 1.86 | 0.66 |
| | H | 1.83 | 0.00 | 2.25 | 2.18 | 0.00 | 1.56 | 82.25 | 1.17 | 5.62 | 0.00 | 0.64 | 0.00 | 2.50 | 0.00 |

| | | | | | | | | | | | | | | | |
|---|---|---|---|---|---|---|---|---|---|---|---|---|---|---|---|
| M08 (4x) | Mean | 3.42 | 2.42 | 2.12 | 3.10 | 1.86 | 1.52 | 69.34 | 3.51 | 1.54 | 2.92 | 1.09 | 3.22 | 3.97 | 0.00 |
| | F | 3.48 | 3.07 | 1.85 | 2.89 | 0.61 | 1.43 | 69.09 | 4.03 | 1.50 | 4.71 | 0.94 | 2.91 | 3.49 | 0.00 |
| | H | 3.35 | 1.76 | 2.39 | 3.31 | 3.11 | 1.61 | 69.60 | 2.98 | 1.57 | 1.13 | 1.23 | 3.52 | 4.45 | 0.00 |
| M09 (4x) | Mean | 1.87 | 1.21 | 1.81 | 1.61 | 1.49 | 1.22 | 35.55 | 1.14 | 29.46 | 0.56 | 0.57 | 3.13 | 1.86 | 18.50 |
| | F | 1.49 | 0.87 | 1.73 | 1.36 | 1.17 | 1.36 | 35.27 | 1.49 | 37.36 | 0.45 | 0.58 | 2.54 | 1.79 | 12.55 |
| | H | 2.24 | 1.56 | 1.89 | 1.87 | 1.82 | 1.09 | 35.84 | 0.80 | 21.57 | 0.67 | 0.57 | 3.71 | 1.93 | 24.46 |
| M10 (2x) | Mean | 2.12 | 2.13 | 1.31 | 2.01 | 0.44 | 0.57 | 77.17 | 0.65 | 2.30 | 2.62 | 0.92 | 5.04 | 2.72 | 0.00 |
| | F | 2.22 | 2.00 | 1.40 | 2.10 | 0.32 | 0.60 | 77.64 | 0.71 | 1.92 | 3.06 | 0.84 | 4.28 | 2.92 | 0.00 |
| | H | 2.03 | 2.25 | 1.22 | 1.92 | 0.56 | 0.55 | 76.70 | 0.59 | 2.68 | 2.18 | 1.00 | 5.80 | 2.52 | 0.00 |
| M11 (2x) | Mean | 2.75 | 3.55 | 1.27 | 2.28 | 0.65 | 0.75 | 66.98 | 0.94 | 1.37 | 2.67 | 0.61 | 12.43 | 3.76 | 0.00 |
| | F | 1.97 | 2.20 | 1.32 | 2.07 | 0.67 | 0.77 | 70.62 | 1.04 | 2.13 | 3.29 | 0.66 | 8.83 | 4.43 | 0.00 |
| | H | 3.53 | 4.89 | 1.22 | 2.48 | 0.63 | 0.72 | 63.35 | 0.84 | 0.62 | 2.05 | 0.55 | 16.02 | 3.09 | 0.00 |
| M12 (4x) | Mean | 1.91 | 0.10 | 1.78 | 2.57 | 0.78 | 2.69 | 82.19 | 0.89 | 1.35 | 0.00 | 0.92 | 0.42 | 4.28 | 0.14 |
| | F | 2.06 | 0.00 | 1.92 | 2.78 | 0.82 | 2.17 | 82.30 | 0.85 | 1.38 | 0.00 | 0.91 | 0.24 | 4.35 | 0.22 |
| | H | 1.75 | 0.19 | 1.64 | 2.36 | 0.74 | 3.21 | 82.07 | 0.92 | 1.32 | 0.00 | 0.92 | 0.60 | 4.21 | 0.06 |
| M13 (4x) | Mean | 1.32 | 0.00 | 0.98 | 1.73 | 0.57 | 0.53 | 69.65 | 1.65 | 19.26 | 0.00 | 0.85 | 0.20 | 2.75 | 0.52 |
| | F | 1.26 | 0.00 | 0.90 | 1.66 | 0.59 | 0.38 | 77.15 | 2.01 | 12.25 | 0.00 | 0.93 | 0.14 | 2.41 | 0.31 |
| | H | 1.38 | 0.00 | 1.06 | 1.80 | 0.54 | 0.69 | 62.14 | 1.29 | 26.26 | 0.00 | 0.77 | 0.26 | 3.08 | 0.72 |
| M14 (2x) | Mean | 2.69 | 0.00 | 2.51 | 3.60 | 0.94 | 0.73 | 82.62 | 0.80 | 0.49 | 0.00 | 0.97 | 0.00 | 4.65 | 0.00 |
| | F | 2.79 | 0.00 | 2.64 | 3.78 | 1.04 | 0.75 | 82.02 | 0.80 | 0.46 | 0.00 | 0.98 | 0.00 | 4.74 | 0.00 |
| | H | 2.59 | 0.00 | 2.38 | 3.42 | 0.85 | 0.71 | 83.21 | 0.81 | 0.52 | 0.00 | 0.96 | 0.00 | 4.55 | 0.00 |
| M15 (4x) | Mean | 1.47 | 0.00 | 1.51 | 2.16 | 0.57 | 1.21 | 86.13 | 0.73 | 1.45 | 0.00 | 1.00 | 0.21 | 3.57 | 0.00 |
| | F | 1.11 | 0.00 | 1.33 | 1.81 | 0.57 | 1.53 | 86.81 | 0.77 | 1.27 | 0.00 | 0.91 | 0.24 | 3.65 | 0.00 |

| | | | | | | | | | | | | | | | |
|---|---|---|---|---|---|---|---|---|---|---|---|---|---|---|---|
| | H | 1.83 | 0.00 | 1.68 | 2.51 | 0.57 | 0.89 | 85.45 | 0.70 | 1.63 | 0.00 | 1.09 | 0.17 | 3.48 | 0.00 |
| M16 (4x) | Mean | 1.21 | 0.10 | 0.96 | 1.67 | 0.52 | 1.90 | 89.49 | 0.75 | 0.54 | 0.00 | 0.53 | 0.46 | 1.88 | 0.00 |
| | F | 1.33 | 0.20 | 1.11 | 1.81 | 0.56 | 2.42 | 86.86 | 0.85 | 0.56 | 0.00 | 0.66 | 0.92 | 2.71 | 0.00 |
| | H | 1.10 | 0.00 | 0.81 | 1.52 | 0.48 | 1.37 | 92.11 | 0.65 | 0.51 | 0.00 | 0.39 | 0.00 | 1.04 | 0.00 |
| M18 (4x) | Mean | 1.18 | 0.00 | 1.06 | 1.68 | 0.58 | 1.72 | 81.81 | 0.93 | 6.01 | 0.00 | 0.86 | 0.31 | 2.94 | 0.93 |
| | F | 1.25 | 0.00 | 1.06 | 1.70 | 0.63 | 2.13 | 84.15 | 1.09 | 3.35 | 0.00 | 0.88 | 0.21 | 2.78 | 0.77 |
| | H | 1.10 | 0.00 | 1.07 | 1.66 | 0.54 | 1.30 | 79.47 | 0.77 | 8.68 | 0.00 | 0.84 | 0.40 | 3.09 | 1.09 |

**Table S3.** Chemotypic characterization of individuals from 13 hermaphroditic (H) and female (F) pools exhibiting the alternative linalool-rich chemotype. Each individual within a pool is classified according to the relative dominance of 1,8-cineole and/or linalool. Values are expressed as the percentage of total quantified compounds. Chemotypes are defined as follows: '1,8-cineole', 1,8-cineole as the dominant compound and linalool <10%; '1,8-cineole*', 1,8-cineole dominant but linalool ≥10% and still substantially lower than 1,8-cineole; '1,8-cineole and linalool', both compounds occur at similar concentrations, with 1,8-cineole reaching at least 25%; "linalool", linalool clearly dominates, with 1,8-cineole concentrations below 25%. Note that in population D07, only individuals from the hermaphroditic pool were analyzed, as the female pool showed no evidence of population-level polymorphism. Samples for which some individuals could not be analyzed are indicated as "not evaluated".

| | **A01-H pool** | **A04-H pool** | **A11-H pool** | **D02-H pool** | **D07-H pool** | **M09-H pool** | **M13-H pool** |
|---|---|---|---|---|---|---|---|
| *Chemotype* | Linalool | 1,8-cineole* | 1,8-cineole* | 1,8-cineole* | 1,8-cineole* | 1,8-cineole and linalool | 1,8-cineole* |
| *1,8-cineole* | 19.05 | 68.49 | 67.53 | 60.97 | 38.21 | 35.84 | 62.14 |
| *Linalool* | 72.00 | 10.51 | 16.45 | 19.79 | 10.66 | 21.57 | 26.26 |
| *Linalool acetate* | 0.271 | 0 | 0.391 | 0 | 0 | 24.46 | 0.719 |

| | **A01-H ind.1** | **A04-H ind.1** | **A11-H ind.1** | **D02-H ind.1** | **D07-H ind.1** | **M09-H ind.1** | **M13-H ind.1** |
|---|---|---|---|---|---|---|---|
| *Chemotype* | Linalool | 1,8-cineole | 1,8-cineole* | 1,8-cineole | Not evaluated | 1,8-cineole | 1,8-cineole and linalool |
| *1,8-cineole* | 23.56 | 87.19 | 72.81 | 81.90 | - | 87.70 | 38.56 |
| *Linalool* | 71.12 | 0.26 | 11.97 | 6.86 | - | 0.70 | 53.75 |
| *Linalool acetate* | 0 | 0 | 0 | 0 | - | 1.07 | 0.81 |

| | **A01-H ind.2** | **A04-H ind.2** | **A11-H ind.2** | **D02-H ind.2** | **D07-H ind.2** | **M09-H ind.2** | **M13-H ind.2** |
|---|---|---|---|---|---|---|---|
| *Chemotype* | Not evaluated | 1,8-cineole | 1,8-cineole* | 1,8-cineole and linalool | 1,8-cineole | 1,8-cineole | 1,8-cineole |
| *1,8-cineole* | - | 82.45 | 71.82 | 43.53 | 54.35 | 89.19 | 83.40 |
| *Linalool* | - | 0.37 | 13.04 | 30.61 | 3.23 | 0.53 | 5.09 |
| *Linalool acetate* | - | 0 | 0.84 | 0 | 0.35 | 0 | 0.70 |

| | **A01-H ind.3** | **A04-H ind.3** | **A11-H ind.3** | **D02-H ind.3** | **D07-H ind.3** | **M09-H ind.3** | **M13-H ind.3** |
|---|---|---|---|---|---|---|---|
| *Chemotype* | 1,8-cineole | 1,8-cineole and linalool | 1,8-cineole* | 1,8-cineole and linalool | Not evaluated | 1,8-cineole | 1,8-cineole |
| *1,8-cineole* | 77.06 | 40.64 | 60.10 | 32.36 | - | 77.0 | 90.50 |
| *Linalool* | 1.27 | 45.17 | 20.50 | 48.0 | - | 0.98 | 1.03 |
| *Linalool acetate* | 0 | 0 | 0.68 | 0 | - | 0 | 0 |

| | **A01-H ind.4** | **A04-H ind.4** | **A11-H ind.4** | **D02-H ind.4** | **D07-H ind.4** | **M09-H ind.4** | **M13-H ind.4** |
|---|---|---|---|---|---|---|---|

| | | | | | | | |
|---|---|---|---|---|---|---|---|
| *Chemotype* | Linalool | 1,8-cineole | 1,8-cineole* | 1,8-cineole* | Not evaluated | Linalool | 1,8-cineole* |
| *1,8-cineole* | 16.79 | 89.68 | 57.97 | 58.47 | - | 8.06 | 66.28 |
| *Linalool* | 76.26 | 0.48 | 27.38 | 15.90 | - | 32.72 | 23.97 |
| *Linalool acetate* | 1.76 | 0 | 1.22 | 0 | - | 44.58 | 0 |

| | **A01-H ind.5** | **A04-H ind.5** | **A11-H ind.5** | **D02-H ind.5** | **D07-H ind.5** | **M09-H ind.5** | **M13-H ind.5** |
|---|---|---|---|---|---|---|---|
| *Chemotype* | Not evaluated | 1,8-cineole | 1,8-cineole | 1,8-cineole | Not evaluated | Linalool | 1,8-cineole* |
| *1,8-cineole* | - | 67.16 | 86.79 | 90.46 | - | 14.96 | 65.84 |
| *Linalool* | - | 8.23 | 2.68 | 2.07 | - | 33.20 | 22.80 |
| *Linalool acetate* | - | 0 | 0.79 | 0 | - | 48.80 | 0 |

| | **A01-F pool** | **A04-F pool** | **A11-F pool** | **D02-F pool** | **D07-F pool** | **M09-F pool** | **M13-F pool** |
|---|---|---|---|---|---|---|---|
| *Chemotype* | 1,8-cineole* | 1,8-cineole* | 1,8-cineole* | 1,8-cineole* | Not evaluated | 1,8-cineole and linalool | 1,8-cineole* |
| *1,8-cineole* | 64.13 | 65.98 | 67.71 | 60.25 | - | 35.27 | 77.15 |
| *Linalool* | 13.69 | 12.78 | 18.51 | 19.37 | - | 37.36 | 12.25 |
| *Linalool acetate* | 0 | 0.49 | 0.10 | 0 | - | 12.55 | 0.314 |

| | **A01-F ind.1** | **A04-F ind.1** | **A11-F ind.1** | **D02-F ind.1** | **D07-F ind.1** | **M09-F ind.1** | **M13-F ind.1** |
|---|---|---|---|---|---|---|---|
| *Chemotype* | 1,8-cineole and linalool | 1,8-cineole | 1,8-cineole* | 1,8-cineole | Not evaluated | Linalool | 1,8-cineole |
| *1,8-cineole* | 54.77 | 84.56 | 69.09 | 60.0 | - | 20.23 | 88.90 |
| *Linalool* | 37.15 | 1.04 | 16.67 | 14.16 | - | 46.74 | 1.36 |
| *Linalool acetate* | 1.09 | 0 | 0 | 0 | - | 27.29 | 0 |

| | **A01-F ind.2** | **A04-F ind.2** | **A11-F ind.2** | **D02-F ind.2** | **D07-F ind.2** | **M09-F ind.2** | **M13-F ind.2** |
|---|---|---|---|---|---|---|---|
| *Chemotype* | Not evaluated | 1,8-cineole | 1,8-cineole and linalool | 1,8-cineole* | Not evaluated | 1,8-cineole | 1,8-cineole* |
| *1,8-cineole* | - | 82.91 | 31.25 | 57.60 | - | 83.63 | 75.48 |
| *Linalool* | - | 0.27 | 62.25 | 30.59 | - | 2.09 | 13.77 |
| *Linalool acetate* | - | 0 | 0.46 | 0 | - | 0 | 0.77 |

| | **A01-F ind.3** | **A04-F ind.3** | **A11-F ind.3** | **D02-F ind.3** | **D07-F ind.3** | **M09-F ind.3** | **M13-F ind.3** |
|---|---|---|---|---|---|---|---|
| *Chemotype* | 1,8-cineole | 1,8-cineole | 1,8-cineole | 1,8-cineole | Not evaluated | Linalool | 1,8-cineole* |
| *1,8-cineole* | 77.15 | 86.23 | 78.38 | 82.0 | - | 15.44 | 69.05 |
| *Linalool* | 0.23 | 1.79 | 3.46 | 2.39 | - | 60.96 | 20.42 |
| *Linalool acetate* | 0 | 0 | 0 | 0 | - | 20.32 | 0.93 |

| | **A01-F ind.4** | **A04-F ind.4** | **A11-F ind.4** | **D02-F ind.4** | **D07-F ind.4** | **M09-F ind.4** | **M13-F ind.4** |
|---|---|---|---|---|---|---|---|
| *Chemotype* | Not evaluated | 1,8-cineole | 1,8-cineole* | 1,8-cineole | Not evaluated | 1,8-cineole | 1,8-cineole* |
| *1,8-cineole* | - | 78.85 | 70.74 | 70.20 | - | 72.42 | 56.90 |
| *Linalool* | - | 0.27 | 11.40 | 0.29 | - | 0.61 | 27.67 |
| *Linalool acetate* | - | 0 | 0.42 | 0 | - | 0 | 6.52 |

| | **A01-F ind.5** | **A04-F ind.5** | **A11-F ind.5** | **D02-F ind.5** | **D07-F ind.5** | **M09-F ind.5** | **M13-F ind.5** |
|---|---|---|---|---|---|---|---|

| | | | | | | | |
|---|---|---|---|---|---|---|---|
| *Chemotype* | 1,8-cineole | 1,8-cineole and linalool | 1,8-cineole | 1,8-cineole | Not evaluated | 1,8-cineole and linalool | 1,8-cineole* |
| *1,8-cineole* | 83.86 | 43.20 | 88.78 | 66.32 | - | 38.64 | 77.69 |
| *Linalool* | 0.54 | 38.41 | 0.38 | 4.15 | - | 19.74 | 13.32 |
| *Linalool acetate* | 0 | 1.89 | 0 | 0 | - | 32.48 | 0.87 |

**Table S4.** Spearman correlation coefficients between the relative abundance of essential oil compounds at the population level and environmental variables. Only significant correlations (adjusted p-value < 0.05 after Bonferroni correction) are highlighted in bold.

| **Compound** | **Environmental variable** | $r_s$ | **p-value** | **Adjusted p-value** |
|---|---|---|---|---|
| 1,8-cineole | pH | 0.103 | 0.384 | 1 |
| 1,8-cineole | Conductivity | 0.057 | 0.629 | 1 |
| 1,8-cineole | OM | 0.067 | 0.571 | 1 |
| 1,8-cineole | WRP | 0.217 | 0.064 | 1 |
| 1,8-cineole | WRC | 0.125 | 0.289 | 1 |
| 1,8-cineole | Bio1 | -0.400 | 0.000 | **0** |
| 1,8-cineole | Bio5 | -0.040 | 0.406 | 1 |
| 1,8-cineole | Bio10 | -0.203 | 0.008 | 0.874 |
| 1,8-cineole | Bio12 | 0.271 | 0.046 | 1 |
| 1,8-cineole | Bio14 | 0.453 | 0.000 | **0** |
| 1,8-cineole | Bio17 | 0.255 | 0.001 | 0.135 |
| 1,8-cineole | MRH | -0.054 | 0.751 | 1 |
| 1,8-cineole | AI | 0.092 | 0.524 | 1 |
| Linalool | pH | 0.071 | 0.547 | 1 |
| Linalool | Conductivity | 0.390 | 0.001 | 0.062 |
| Linalool | OM | 0.406 | 0.000 | **0.031** |
| Linalool | WRP | 0.362 | 0.002 | 0.166 |
| Linalool | WRC | 0.392 | 0.001 | 0.062 |
| Linalool | Bio1 | -0.020 | 0.920 | 1 |
| Linalool | Bio5 | 0.322 | 0.002 | 0.229 |
| Linalool | Bio10 | 0.204 | 0.041 | 1 |
| Linalool | Bio12 | -0.024 | 0.877 | 1 |
| Linalool | Bio14 | 0.204 | 0.183 | 1 |
| Linalool | Bio17 | 0.295 | 0.040 | 1 |
| Linalool | MRH | -0.525 | 0.000 | **0** |
| Linalool | AI | -0.295 | 0.018 | 1 |
| Linalool acetate | pH | 0.329 | 0.004 | 0.437 |
| Linalool acetate | Conductivity | 0.311 | 0.007 | 0.738 |
| Linalool acetate | OM | 0.231 | 0.048 | 1 |
| Linalool acetate | WRP | 0.270 | 0.020 | 1 |
| Linalool acetate | WRC | 0.261 | 0.025 | 1 |

| | | | | |
|---|---|---|---|---|
| Linalool acetate | Bio1 | -0.366 | 0.003 | 0.291 |
| Linalool acetate | Bio5 | -0.054 | 0.782 | 1 |
| Linalool acetate | Bio10 | -0.226 | 0.105 | 1 |
| Linalool acetate | Bio12 | 0.046 | 0.767 | 1 |
| Linalool acetate | Bio14 | 0.265 | 0.073 | 1 |
| Linalool acetate | Bio17 | 0.371 | 0.006 | 0.582 |
| Linalool acetate | MRH | -0.324 | 0.003 | 0.281 |
| Linalool acetate | AI | -0.079 | 0.486 | 1 |
| Camphor | pH | -0.001 | 0.995 | 1 |
| Camphor | Conductivity | -0.397 | 0.001 | 0.052 |
| Camphor | OM | -0.337 | 0.003 | 0.343 |
| Camphor | WRP | -0.330 | 0.004 | 0.426 |
| Camphor | WRC | -0.415 | 0.000 | **0.021** |
| Camphor | Bio1 | 0.522 | 0.000 | **0** |
| Camphor | Bio5 | 0.037 | 0.494 | 1 |
| Camphor | Bio10 | 0.225 | 0.010 | 0.998 |
| Camphor | Bio12 | -0.307 | 0.019 | 1 |
| Camphor | Bio14 | -0.373 | 0.000 | **0.010** |
| Camphor | Bio17 | -0.399 | 0.000 | **0** |
| Camphor | MRH | 0.217 | 0.228 | 1 |
| Camphor | AI | -0.041 | 0.818 | 1 |
| Borneol | pH | -0.149 | 0.205 | 1 |
| Borneol | Conductivity | -0.260 | 0.025 | 1 |
| Borneol | OM | -0.305 | 0.008 | 0.863 |
| Borneol | WRP | -0.452 | 0.000 | **0.010** |
| Borneol | WRC | -0.326 | 0.005 | 0.478 |
| Borneol | Bio1 | 0.359 | 0.000 | **0.021** |
| Borneol | Bio5 | -0.255 | 0.075 | 1 |
| Borneol | Bio10 | -0.034 | 0.634 | 1 |
| Borneol | Bio12 | -0.278 | 0.030 | 1 |
| Borneol | Bio14 | -0.488 | 0.000 | **0** |
| Borneol | Bio17 | -0.597 | 0.000 | **0** |
| Borneol | MRH | 0.404 | 0.005 | 0.499 |
| Borneol | AI | 0.145 | 0.258 | 1 |
| Camphene | pH | -0.107 | 0.363 | 1 |
| Camphene | Conductivity | -0.310 | 0.007 | 0.749 |
| Camphene | OM | -0.327 | 0.004 | 0.458 |
| Camphene | WRP | -0.441 | 0.000 | **0.010** |
| Camphene | WRC | -0.379 | 0.001 | 0.094 |
| Camphene | Bio1 | 0.429 | 0.000 | **0** |
| Camphene | Bio5 | -0.184 | 0.250 | 1 |
| Camphene | Bio10 | 0.028 | 0.315 | 1 |
| Camphene | Bio12 | -0.284 | 0.022 | 1 |

| | | | | |
|---|---|---|---|---|
| Camphene | Bio14 | -0.466 | 0.000 | **0** |
| Camphene | Bio17 | -0.548 | 0.000 | **0** |
| Camphene | MRH | 0.379 | 0.011 | 1 |
| Camphene | AI | 0.119 | 0.388 | 1 |
| α-terpineol | pH | -0.246 | 0.035 | 1 |
| α-terpineol | Conductivity | 0.096 | 0.414 | 1 |
| α-terpineol | OM | 0.066 | 0.579 | 1 |
| α-terpineol | WRP | 0.194 | 0.098 | 1 |
| α-terpineol | WRC | 0.141 | 0.232 | 1 |
| α-terpineol | Bio1 | 0.087 | 0.321 | 1 |
| α-terpineol | Bio5 | 0.203 | 0.082 | 1 |
| α-terpineol | Bio10 | 0.192 | 0.068 | 1 |
| α-terpineol | Bio12 | 0.110 | 0.178 | 1 |
| α-terpineol | Bio14 | 0.064 | 0.872 | 1 |
| α-terpineol | Bio17 | 0.143 | 0.402 | 1 |
| α-terpineol | MRH | 0.020 | 0.907 | 1 |
| α-terpineol | AI | -0.121 | 0.638 | 1 |
| α-pinene | pH | -0.291 | 0.012 | 1 |
| α-pinene | Conductivity | -0.226 | 0.053 | 1 |
| α-pinene | OM | -0.249 | 0.032 | 1 |
| α-pinene | WRP | -0.371 | 0.001 | 0.125 |
| α-pinene | WRC | -0.323 | 0.005 | 0.510 |
| α-pinene | Bio1 | 0.428 | 0.000 | **0** |
| α-pinene | Bio5 | -0.022 | 0.798 | 1 |
| α-pinene | Bio10 | 0.174 | 0.020 | 1 |
| α-pinene | Bio12 | -0.101 | 0.568 | 1 |
| α-pinene | Bio14 | -0.442 | 0.000 | **0** |
| α-pinene | Bio17 | -0.311 | 0.001 | 0.052 |
| α-pinene | MRH | 0.355 | 0.029 | 1 |
| α-pinene | AI | 0.103 | 0.341 | 1 |

**Table S5.** Spearman correlation coefficients between the relative abundance of essential oil compounds at the individual level and environmental variables. Individual-level data include both separately analyzed individuals and those from hermaphroditic and female pools (see **Methods** for details). Only significant correlations (adjusted p-value < 0.05 after Bonferroni correction) are highlighted in bold.

| Compound | Environmental variable | $r_S$ | p-value | Adjusted p-value |
|---|---|---|---|---|
| 1,8-cineole | pH | 0.127 | 0.009 | 0.915 |
| 1,8-cineole | Conductivity | 0.104 | 0.032 | 1 |
| 1,8-cineole | OM | 0.12 | 0.014 | 1 |

| | | | | |
|---|---|---|---|---|
| 1,8-cineole | WRP | 0.205 | 0 | **0** |
| 1,8-cineole | WRC | 0.176 | 0 | **0.031** |
| 1,8-cineole | Bio1 | -0.351 | 0 | **0** |
| 1,8-cineole | Bio5 | 0.011 | 0.37 | 1 |
| 1,8-cineole | Bio10 | -0.144 | 0 | **0** |
| 1,8-cineole | Bio12 | 0.244 | 0 | **0** |
| 1,8-cineole | Bio14 | 0.341 | 0 | **0** |
| 1,8-cineole | Bio17 | 0.262 | 0 | **0** |
| 1,8-cineole | MRH | -0.111 | 0.666 | 1 |
| 1,8-cineole | AI | 0.036 | 0.682 | 1 |
| Linalool | pH | -0.043 | 0.374 | 1 |
| Linalool | Conductivity | 0.34 | 0 | **0** |
| Linalool | OM | 0.367 | 0 | **0** |
| Linalool | WRP | 0.382 | 0 | **0** |
| Linalool | WRC | 0.351 | 0 | **0** |
| Linalool | Bio1 | -0.012 | 0.968 | 1 |
| Linalool | Bio5 | 0.312 | 0 | **0** |
| Linalool | Bio10 | 0.199 | 0 | **0** |
| Linalool | Bio12 | -0.054 | 0.532 | 1 |
| Linalool | Bio14 | 0.289 | 0 | **0** |
| Linalool | Bio17 | 0.266 | 0 | **0** |
| Linalool | MRH | -0.473 | 0 | **0** |
| Linalool | AI | -0.283 | 0 | **0** |
| Linalool acetate | pH | 0.284 | 0 | **0** |
| Linalool acetate | Conductivity | 0.146 | 0.003 | 0.27 |
| Linalool acetate | OM | 0.09 | 0.064 | 1 |
| Linalool acetate | WRP | 0.11 | 0.024 | 1 |
| Linalool acetate | WRC | 0.113 | 0.02 | 1 |
| Linalool acetate | Bio1 | -0.283 | 0 | **0** |
| Linalool acetate | Bio5 | -0.109 | 0.046 | 1 |
| Linalool acetate | Bio10 | -0.242 | 0 | **0** |
| Linalool acetate | Bio12 | -0.005 | 0.852 | 1 |
| Linalool acetate | Bio14 | 0.215 | 0.001 | 0.052 |
| Linalool acetate | Bio17 | 0.171 | 0.005 | 0.562 |
| Linalool acetate | MRH | -0.149 | 0.001 | 0.052 |
| Linalool acetate | AI | -0.031 | 0.489 | 1 |
| Camphor | pH | -0.012 | 0.808 | 1 |
| Camphor | Conductivity | -0.348 | 0 | **0** |
| Camphor | OM | -0.321 | 0 | **0** |
| Camphor | WRP | -0.269 | 0 | **0** |
| Camphor | WRC | -0.394 | 0 | **0** |
| Camphor | Bio1 | 0.437 | 0 | **0** |
| Camphor | Bio5 | -0.042 | 0.982 | 1 |

| | | | | |
|---|---|---|---|---|
| Camphor | Bio10 | 0.134 | 0 | **0** |
| Camphor | Bio12 | -0.3 | 0 | **0** |
| Camphor | Bio14 | -0.322 | 0 | **0** |
| Camphor | Bio17 | -0.394 | 0 | **0** |
| Camphor | MRH | 0.228 | 0.002 | 0.156 |
| Camphor | AI | -0.006 | 0.937 | 1 |
| Borneol | pH | -0.182 | 0 | **0.021** |
| Borneol | Conductivity | -0.25 | 0 | **0** |
| Borneol | OM | -0.305 | 0 | **0** |
| Borneol | WRP | -0.387 | 0 | **0** |
| Borneol | WRC | -0.331 | 0 | **0** |
| Borneol | Bio1 | 0.302 | 0 | **0** |
| Borneol | Bio5 | -0.233 | 0 | **0** |
| Borneol | Bio10 | -0.039 | 0.422 | 1 |
| Borneol | Bio12 | -0.28 | 0 | **0** |
| Borneol | Bio14 | -0.379 | 0 | **0** |
| Borneol | Bio17 | -0.538 | 0 | **0** |
| Borneol | MRH | 0.367 | 0 | **0** |
| Borneol | AI | 0.117 | 0.023 | 1 |
| Camphene | pH | -0.146 | 0.003 | 0.26 |
| Camphene | Conductivity | -0.262 | 0 | **0** |
| Camphene | OM | -0.29 | 0 | **0** |
| Camphene | WRP | -0.341 | 0 | **0** |
| Camphene | WRC | -0.349 | 0 | **0** |
| Camphene | Bio1 | 0.364 | 0 | **0** |
| Camphene | Bio5 | -0.19 | 0.003 | 0.312 |
| Camphene | Bio10 | -0.001 | 0.084 | 1 |
| Camphene | Bio12 | -0.287 | 0 | **0** |
| Camphene | Bio14 | -0.386 | 0 | **0** |
| Camphene | Bio17 | -0.482 | 0 | **0** |
| Camphene | MRH | 0.334 | 0 | **0** |
| Camphene | AI | 0.1 | 0.088 | 1 |
| α-terpineol | pH | -0.2 | 0 | **0** |
| α-terpineol | Conductivity | 0.042 | 0.386 | 1 |
| α-terpineol | OM | 0.053 | 0.281 | 1 |
| α-terpineol | WRP | 0.12 | 0.013 | 1 |
| α-terpineol | WRC | 0.126 | 0.009 | 0.967 |
| α-terpineol | Bio1 | 0.078 | 0.021 | 1 |
| α-terpineol | Bio5 | 0.144 | 0.003 | 0.281 |
| α-terpineol | Bio10 | 0.152 | 0 | **0.031** |
| α-terpineol | Bio12 | 0.124 | 0.001 | 0.114 |
| α-terpineol | Bio14 | 0.001 | 0.381 | 1 |

| | | | | |
|---|---|---|---|---|
| **α**-terpineol | Bio17 | 0.085 | 0.42 | 1 |
| **α**-terpineol | MRH | 0.062 | 0.273 | 1 |
| **α**-terpineol | AI | -0.048 | 0.968 | 1 |
| **α**-pinene | pH | -0.303 | 0 | **0** |
| **α**-pinene | Conductivity | -0.154 | 0.002 | 0.156 |
| **α**-pinene | OM | -0.159 | 0.001 | 0.104 |
| **α**-pinene | WRP | -0.305 | 0 | **0** |
| **α**-pinene | WRC | -0.21 | 0 | **0** |
| **α**-pinene | Bio1 | 0.37 | 0 | **0** |
| **α**-pinene | Bio5 | -0.031 | 0.651 | 1 |
| **α**-pinene | Bio10 | 0.146 | 0 | **0** |
| **α**-pinene | Bio12 | -0.103 | 0.134 | 1 |
| **α**-pinene | Bio14 | -0.401 | 0 | **0** |
| **α**-pinene | Bio17 | -0.243 | 0 | **0** |
| **α**-pinene | MRH | 0.308 | 0 | **0** |
| **α**-pinene | AI | 0.09 | 0.044 | 1 |

Figure S1

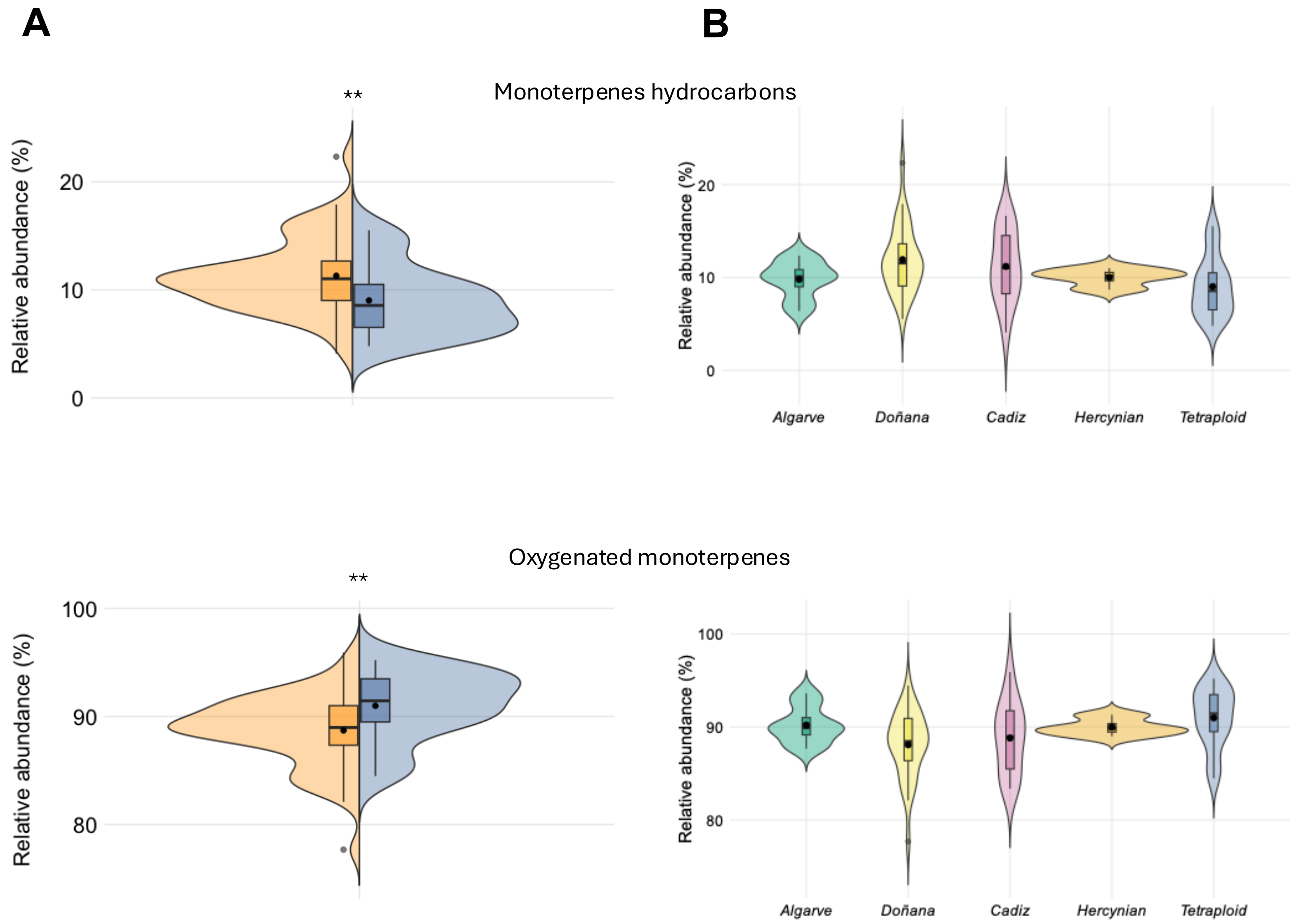

A
B
Monoterpenes hydrocarbons
**
Relative abundance (%)
20
10
0
Algarve
Doñana
Cadiz
Hercynian
Tetraploid
Oxygenated monoterpenes
**
Relative abundance (%)
100
90
80
Algarve
Doñana
Cadiz
Hercynian
Tetraploid
